# Optimizing stability in dynamic small-molecule binding proteins

**DOI:** 10.1101/2025.07.07.663574

**Authors:** Marc Scherer, Mark Kriegel, Birte Höcker, Sarel J. Fleishman

**Affiliations:** University of Bayreuth, Department of Biochemistry, Bayreuth, Germany; Weizmann Institute of Science, Department of Biomolecular Sciences, Rehovot, Israel

**Keywords:** protein dynamics, computational design, PROSS, Gaussian network models, ligand binding

## Abstract

The function of dynamic proteins is determined by the stability of distinct conformational states and the energy barriers that separate these states. For most dynamic proteins, the molecular details of the energy barriers are not known, implying a fundamental limit to the ability of protein design methods to engineer beneficial mutations without disrupting activity. We hypothesized that designing mutations that are compatible with structurally distinct equilibrium conformations may enable reliable stability design. We focus on periplasmic binding proteins (PBPs), a superfamily of dynamic proteins that change conformation from open to closed states in response to binding their small-molecule ligands. We find that the evolutionary constrained space of allowed mutations computed for one conformation is incompatible with the other. Therefore, putative conformational hinge points and interface residues were additionally constraint, and incompatible mutations were filtered out. Starting from four different PBPs, we designed a total of 16 stabilized variants with 7-28 mutations each. Our results show that design based on a single conformation with evolutionary constraints is not sufficient to maintain wild type-like binding affinity. Conversely, using a subset of mutations compatible with both conformations and structural constraints reliably enhances thermal stability while mitigating trade-offs in ligand binding. Our work demonstrates a straightforward method for the one-shot stabilization of dynamic proteins, which is critically required to generate robust starting points for thermostable and responsive biosensors.

## Introduction

Many proteins are dynamic, alternating between several conformational states in equilibrium or in response to binding other molecules.^1,2^ Protein dynamics enable transitions between functionally different states, allowing for complex outcomes such as allosteric regulation and cooperativity.^3^ The folding landscapes of dynamic proteins may have been tuned by evolution to facilitate rapid (nanosecond-to-second) exchange between functional states while preventing the protein from being locked in a single conformation.^4–6^ Notwithstanding their importance, the energy landscapes governing protein conformational equilibria are complex, remain poorly understood, and are difficult to estimate computationally, except in the case of small, fast-folding proteins.^7–9^

Our paper focuses on improving the stability of dynamic binding proteins. In the case of proteins that exhibit limited dynamics, reliable methods that combine phylogenetic analysis and energy calculations applied to a single conformation, such as PROSS (protein repair one-stop shop) and FireProt, have shown high reliability and broad scope.^10–13^ When applied to dynamic proteins, however, the results of stability design have been mixed. For instance, PROSS has been successfully applied to improve the stability, expressibility and tendency to form crystals of human estrogen receptor ɑ (hERɑ) based on a single ligand- and coactivator-bound crystal structure^14^ and to improve the expressibility of a voltage-gated potassium (Kv) channel;^15^ in both cases, without a discernible drop in activity. Notably, inter-subunit interfaces and putative hinge points were restricted from mutation in both cases. In contrast, applying PROSS to the periplasmic binding protein (PBP) DalS^16^ and the amino ester hydrolase QVH^17^ (without considering hinge points) decreased activity. PBPs may be an exceptionally challenging target for computational design as a previous design study^18^ showed unpredicted conformational changes, loss of stability, and undesired functional profiles.^19^

Bacterial PBPs are a diverse superfamily responsible for capturing and translocating small molecules from the bacterial periplasm, across the plasma membrane, into the cytosol.^20–22^ We focus this study on PBPs because of the general interest in improving the stability of dynamic proteins, the specific difficulties observed in previous design studies^18,19^ and the many potential applications in biosensing of these proteins.^23^ PBPs share a two-lobe architecture connected by a hinge region and employ a ‘Venus flytrap’ mechanism switching from an open to a closed conformation to bind various ligand molecules.^6,24,25^ Despite a shared structural fold, PBPs possess diverse sequences,^20,26^ and their energy landscapes are complex, featuring multiple meta-stable states.^27,28^ Importantly, strategies were developed to convert PBPs into fluorescence-based biosensors primarily for biomedical research. ^29–32^ But, inserting fluorescent protein domains or optimizing the ligand specificity and binding affinity of a biosensor are frequently accompanied by reduced expression yields ^33^ and reduced thermal stability.^16^ A reliable strategy for stabilizing PBPs may therefore be particularly helpful for biosensor engineering.

We show that restricting stabilizing mutations to those that are compatible with both the open and closed states and away from putative hinge points is an effective way to design stable PBPs. We thus provide a strategy to optimize stability in dynamic proteins and obtain suitable starting points for the engineering of PBP-based biosensors.

## Results

### Conformation impacts stability design choices

We began by searching structural databases for PBPs that undergo large conformational changes upon ligand binding and have been structurally characterized in both open and closed states. We selected four PBPs (PotF, TphC, MBP, and LAO) that bind to diverse ligands, ranging from aromatic dicarboxylic acids to sugars (Figure 1A; PDB IDs can be found in Table S1). To serve as a reference, we applied PROSS design calculations with standard parameters on the open and closed structures of each PBP (Figure 1B, Design X.1 & X.2; see Methods for details).

**Figure 1.**
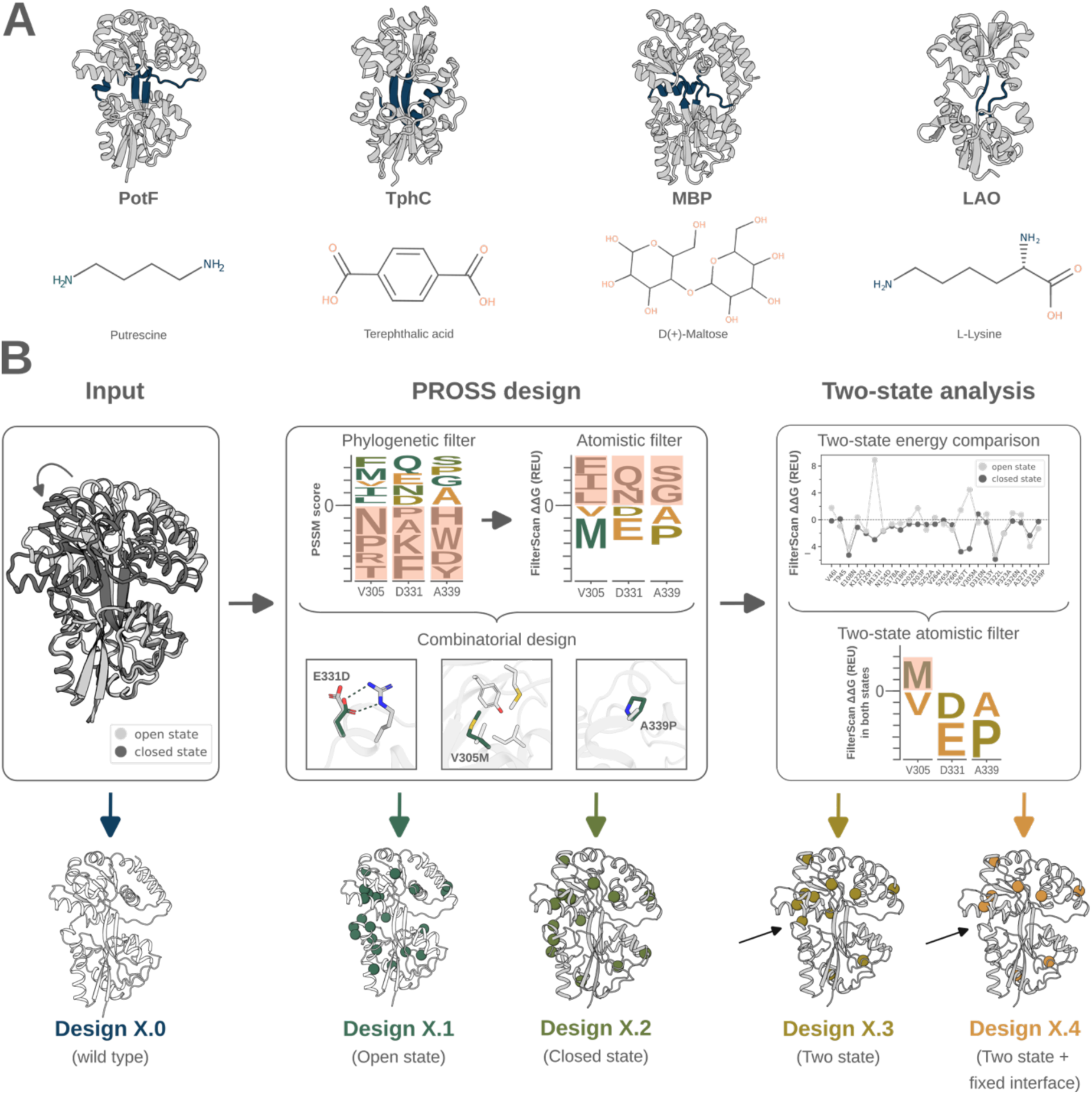
Design strategy for optimizing PBP stability with single- and two-state PROSS design. (A) Four PBPs were chosen: putrescine-binding protein PotF; terephthalic acid-binding protein TphC; maltose-binding protein MBP; and lysine-binding protein LAO. Closed-state crystal structures (gray) with hinge regions (blue) are shown as cartoons. (B) The design strategy used the open- and closed-state crystal structures of PotF, TphC, MBP and LAO as input. The wild type sequences of all PBPs were used as control (Designs X.0). PROSS design calculations were run with default parameters, generating Designs X.1 and X.2 based on the open and closed structures, respectively. In the inset, three PROSS mutations of PotF showcase the formation of novel hydrogen bonds, improved core packing and loop rigidification. The impact of mutations on the energy (ΔΔ*G*) depends on the backbone conformation between the two states. Hence, mutations that were destabilizing (positive energies) in any state were filtered out leading to Design X.3. Design X.4 was generated by fixing the hinge region and lobe interface before running PROSS design calculations and the two-state energy filter. The black arrows in Design X.3-4 indicate one mutation that was not present in Design X.4 due to the fixed hinge region.

We compared the ΔΔ*G* values for all mutations of the open and closed-state designs. This analysis showed that some mutations were stabilizing in one and destabilizing in the other state (Figure 2A). Generally, the open-state energies tended to be lower (more favorable) than the closed-state ones, likely due to the high packing density in the latter. Inspection of the residues around PROSS mutations that showed state-dependent sign differences in energy indicated subtle differences in backbone and side-chain configurations (Figure 2B). We classified the mutated positions into surface, boundary and core layers, finding that the energies of surface mutations were similar in both states, likely reflecting reduced sensitivity due to low packing density (Figure S1). By contrast, boundary and core mutations, especially in PotF and TphC, showed sign energy differences. We concluded that mutations in the core of the protein are especially sensitive to the conformational state.

**Figure 2.**
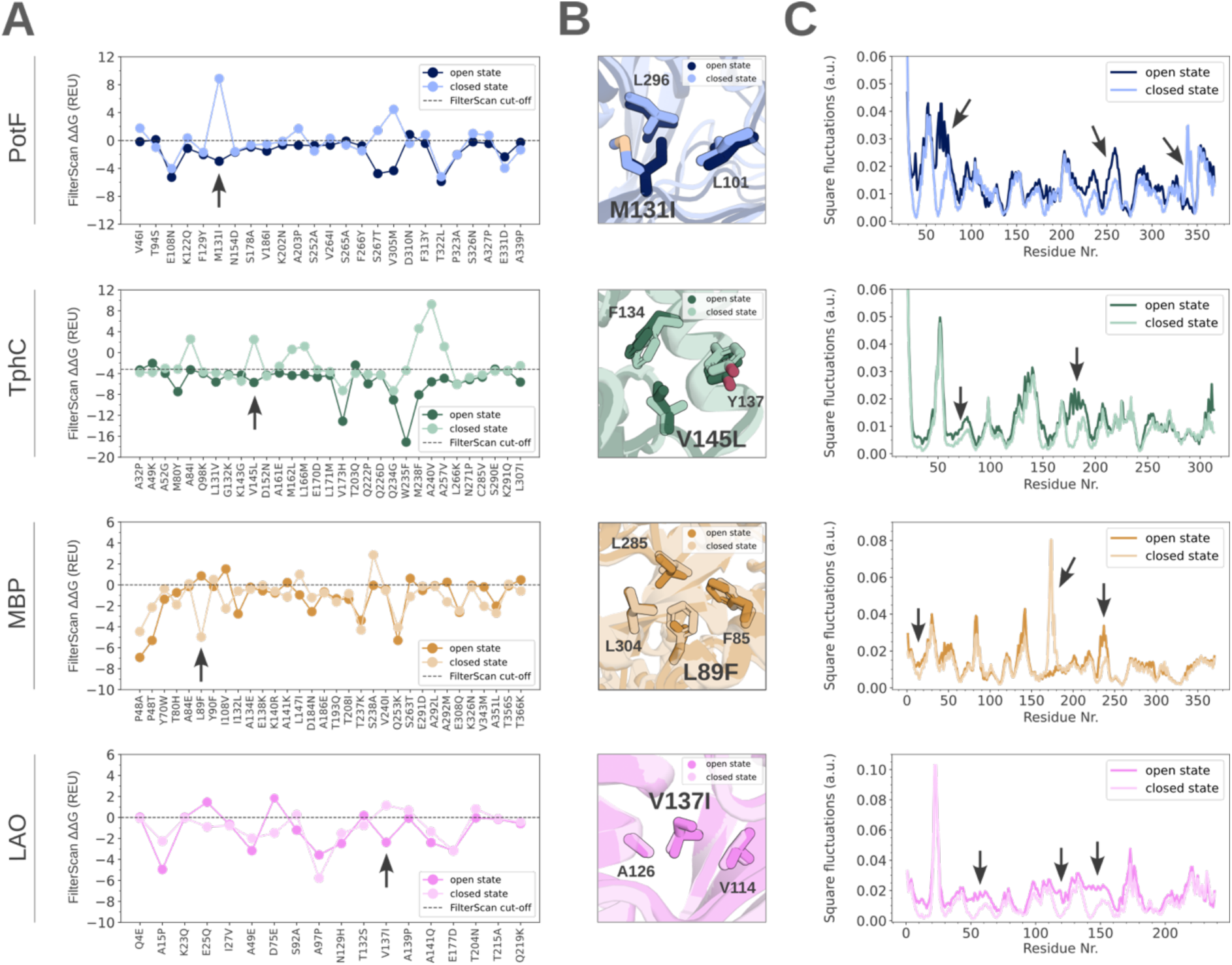
Rosetta ΔΔ*G* and packing density analysis of PBP single-state PROSS mutations. (A) ΔΔ*G* values of PROSS mutations generated based on the open and closed state of PotF (blue), TphC (green), MBP (orange), and LAO (pink), and cut-off energies (dotted lines). Energies are plotted in Rosetta energy units (REU). Arrows indicate mutations visualized in (B). (B) Subtle packing differences of residues were observed around four representative mutations (shown as sticks) of PotF (blue), TphC (green), MBP (orange) and LAO (pink) designs, respectively. Mutations were shown to have stabilizing ΔΔ*G* values in one state and destabilizing values in the other state as part of the analysis in (A). (C) Backbone square fluctuations (inverse of the packing density) of PotF (blue), TphC (green), MBP (orange) and LAO (pink) based on GNMs. Arrows indicate large packing density changes of hinge and interface residues.

Two additional sets of designs were generated: To avoid changes in the conformational equilibrium of PBPs, mutations that showed state-dependent sign differences in energy were excluded in the third set (Figure 1B, Design X.3). Still, mutations in the interface and hinge region were present. As those areas showed the strongest changes in local backbone density as seen by Gaussian Network Model (GNM) calculations (Figure 2C), in the fourth set of designs we also eliminated mutations in the hinge or lobe interface (Methods; Figure 1B, Figure S2, Design X.4). In total, 16 PBP designs (4 designs per wild type protein) were selected for experimental testing and compared to their respective wild type (WT) (Table 1, Table S2 and Figures S3-6).

**Table 1.** Thermal stability and binding parameters for the PBP designs and wild types.

| Protein target | Design ID | Nr. of mutations | Mutational load | Apparent $T_M$ in °C | $K_D$ (nM) |
| --- | --- | --- | --- | --- | --- |
| PotF | PotF.0 | 0 | 0 % | $63 \pm 0.4$ | $42 \pm 3$ |
| | PotF.1 | 22 | 6 % | $76 \pm 0.3$ | $479 \pm 12$ |
| | PotF.2 | 14 | 4 % | $73 \pm 0.3$ | $42 \pm 13$ |
| | PotF.3 | 12 | 4 % | $73 \pm 0.3$ | $101 \pm 17$ |
| | PotF.4 | 9 | 3 % | $75 \pm 0.6$ | $103 \pm 10$ |
| TphC | TphC.0 | 0 | 0 % | $61 \pm 0.1$ | $1180 \pm 250$ |
| | TphC.1 | 28 | 10 % | $68 \pm 0.0$ | $5750 \pm 709$ |
| | TphC.2 | 20 | 7 % | $68 \pm 0.1$ | $627 \pm 69$ |
| | TphC.3 | 16 | 5 % | $66 \pm 0.1$ | $716 \pm 91$ |
| | TphC.4 | 9 | 3 % | $67 \pm 0.1$ | $282 \pm 39$ |
| MBP | MBP.0 | 0 | 0 % | $63 \pm 0.1$ | $1240 \pm 102$ |
| | MBP.1 | 25 | 7 % | $80 \pm 0.1$ | $1310 \pm 293$ |
| | MBP.2 | 25 | 7 % | $79 \pm 0.1$ | $<10 \mu\text{M}$ |
| | MBP.3 | 18 | 5 % | $78 \pm 0.1$ | $450 \pm 44$ |
| | MBP.4 | 14 | 4 % | $77 \pm 0.2$ | $2960 \pm 675$ |
| LAO | LAO.0 | 0 | 0 % | $47 \pm 0.1$ | $93 \pm 35$ |
| | LAO.1 | 13 | 5 % | $51 \pm 0.1$ | $2496 \pm 211$ |
| | LAO.2 | 13 | 5 % | $55 \pm 0.1$ | $612 \pm 56$ |
| | LAO.3 | 9 | 4 % | $54 \pm 0.1$ | $824 \pm 18$ |
| | LAO.4 | 7 | 3 % | $54 \pm 0.2$ | $150 \pm 21$ |

### Two-state filter and structural constraints enable reliable stability optimization

WT proteins and designs were expressed in the cytosol of *E. coli* and purified. First, the folding of all PBP designs was checked with circular dichroism (CD) spectroscopy (Figure S7). Subsequent thermal melt (Tmelt) experiments revealed that all PBP designs exhibited improved thermal stability with apparent Δ*T_m_* values ranging between 4 - 17 ℃ (Figure 3A and S8, Table 1). For comparison, the apparent *T_m_* values of wild type PotF, TphC, MBP and LAO were 63 ℃, 61 ℃, 63 ℃ and 47 ℃, respectively. Remarkably, in case of PotF, TphC and LAO, the increase in apparent *T_m_* was not correlated with the number of mutations, opposing a trend seen in previous work.^34^

**Figure 3.**
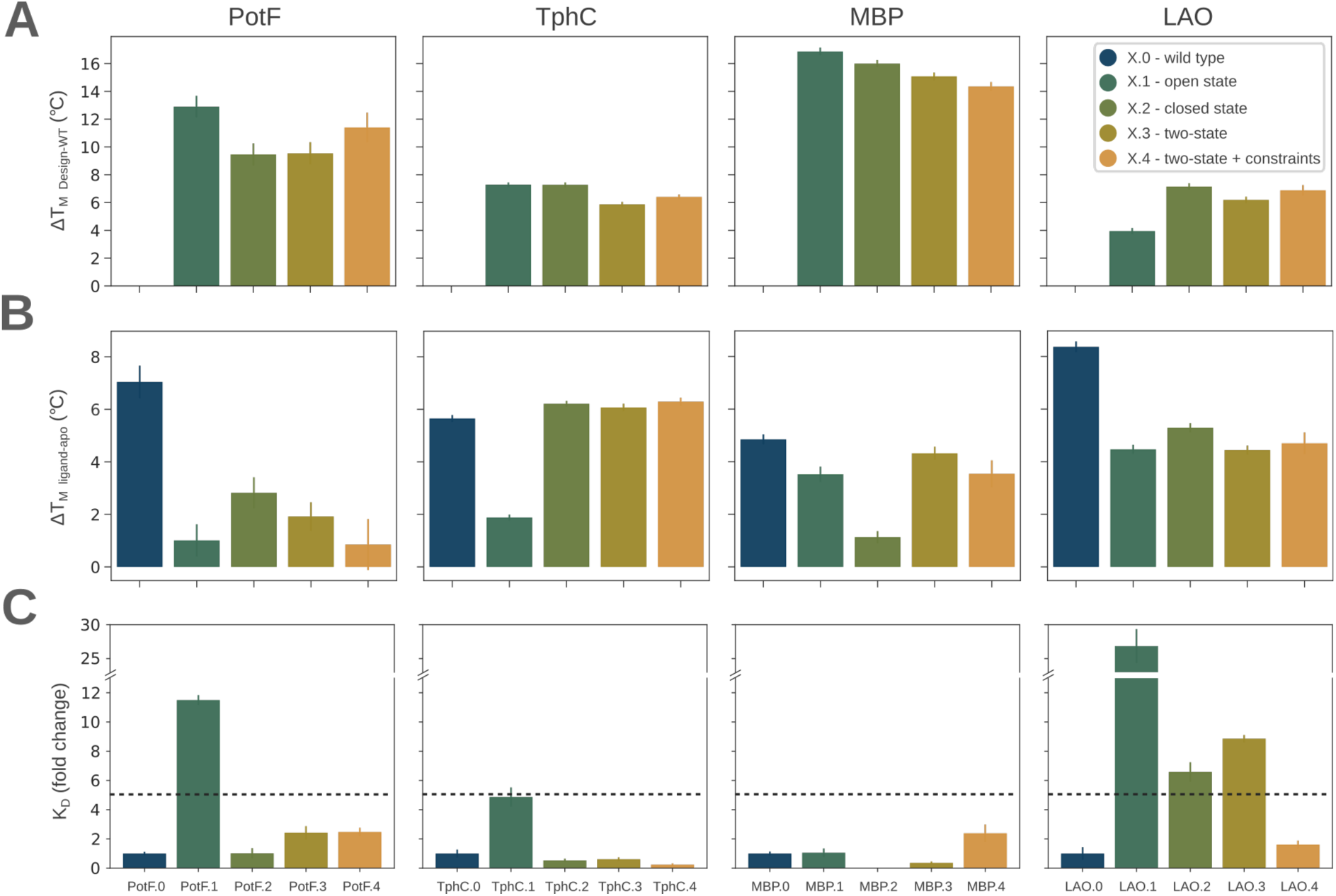
Thermostability and binding affinity of PBP WTs and designs. (A) The change in apparent melting temperature Δ*T_m_* was calculated relative to the WT. (B) The change in the apparent Δ*T_m_* upon addition of 100 µM ligand was calculated by subtracting the respective sample with the apo protein control. (C) Fold change of *K_D_* against WT. Error bars reflect the standard deviation of three independent repeats. Putrescine, terephthalic acid, maltose and L-lysine were used for PotF, TphC, MBP and LAO binding studies, respectively. A fivefold decreased affinity compared to the WT is visualized by a dashed line.

Next, we investigated whether the PROSS designs bind their cognate ligands. Tmelt experiments were performed in the presence of 100 µM of the cognate ligands, putrescine, terephthalic acid, maltose or L-lysine in case of PotF, TphC, MBP, and LAO designs, respectively. Using a threshold of +1 ℃ Δ*T_m_*, all PBP designs bound their cognate ligands but Des1.4 (Table S4, Figure S9). Except for three TphC designs, PBP PROSS designs exhibited slightly lower Δ*T_m_* (ligand-apo) values than the wild-type proteins (Figure 3B). These results indicate that the binding affinity of most PBP variants to their respective ligands might be decreased. To investigate the effect of the mutations on the ligand binding behavior in more detail, isothermal titration calorimetry (ITC) measurements of the PBP designs were conducted and compared with the binding affinities (*K_D_*) of their respective WT (Figure 3C and Figures S10-13). The binding affinities of the PBP WT proteins were determined to 42 ± 3 nM for PotF, 1180 ± 250 nM for TphC, 1240 ± 102 nM for MBP, and 93 ± 35 nM for LAO which were in accordance with previously published binding constants: 68 ± 40 nM for PotF^28^, 364 nM for TphC^35^, 1102 ± 65 nM for MBP^36^, and 232 ± 106 nM for LAO^37^. Three out of eight single-state designs exhibited a decreased affinity of fivefold or more; two of these were designed based on the open state. For the single-state MBP.2, which is based on the closed state, the ITC data could not be properly fitted (see Table 1 and Figures S12). The two-state filtered designs without hinge constraints, on the other hand, showed decreased binding affinity only in one out of four cases, while all two-state filtered designs with hinge constraints showed WT-like affinity. The binding affinity was marginally improved (less than fivefold) for TphC.2-4 and MBP.3. Taken together, these data suggest that designing PBPs from only one state is more likely to change binding affinity. Two-state mutation filtering, especially with constraints on the hinge region, on the other hand, reliably enhanced the thermal stability of all PBPs while retaining their ligand-binding affinity.

## Discussion

Optimizing dynamic proteins without prior knowledge of the underlying energy landscape is a challenging task, as conformational states and function are usually dependent. Our thermal melt and binding data revealed that evolutionary constraints on the sequence space in single-state design of PBPs were not sufficient to maintain WT-like binding affinity. Fine balancing is likely needed to allow PBP open-to-closed transitions in fast timescales (low energy barriers) but also stabilize an open state ensemble for the ligand to enter. The packing density differences between the open and closed state revealed by mutational energy comparison might be involved in maintaining this conformational balance. Mutations in the binding pocket of PBPs can have an effect on the conformational ensemble and accessibility of closed states.^38,39^ Similarly, mutations in the hinge area can impair ligand-binding^36,40^ by differentially stabilizing particular backbone conformations. Our data revealed that even remote mutations can have state-dependent sign differences in energy. Taking advantage of structural information of a second state improved the reliability of the design approach but still led to decreased binding affinity in the case of LAO.3. Additional constraints on residues of the hinge and interface were necessary to achieve success in all cases (Designs X.4) while still improving thermal stability at a similar level as designs with more than twice the number of mutations. Hence, the approach presented here allowed reliable optimization of PBPs at a low mutational load. Importantly, the workflow described here can generate robust starting points for engineering biosensors for which a general stability optimization method is still missing.

While our design approach worked well on four PBPs, it remains to be seen whether the approach generalizes to other dynamic proteins. For most dynamic proteins, high-quality structural information of all functional states is not available. If modelling of structural ensembles of dynamic proteins can be achieved with high precision using machine learning-based approaches such as AlphaFold^41,42^, the applicability of the design approach could be broadened. We saw that at least one out of five models (default setting) generated by AlphaFold3 resembled the open and closed conformations of the PBP wild type structures studied here. Still, it must be ensured that the chosen structural representations constitute functional states of the protein as the stabilization of non-functional conformations will negatively impact function. We provide a design method for the stabilization of dynamic proteins and present initial steps towards understanding and designing complex dynamic proteins.

## Methods

### Sequence design with PROSS

PROSS^10,11^ uses the crystal structure and amino acid sequence of a protein as input. First, a multiple sequence alignment (MSA) is generated that serves as a basis to compute a position-specific scoring matrix (PSSM) reflecting the likelihood of an amino acid to occur at each position of the protein in the natural diversity. Filtering out all amino acid identities that do not occur frequently in the MSA (PSSM values < 0), allows to focus the sequence space on regions that evolution has explored and can be considered more likely to maintain the protein fold. Subsequently, Rosetta atomistic design calculations of each single amino acid substitution identified stabilizing mutations (Rosetta FilterScan functionality calculates ΔΔ*G* values of mutations compared to WT) which were then considered for a final combinatorial design step. For that, nine energy cut-offs (−4, −3.20, −2.88, −2.4, −2, −1.6, −1.2, −0.72, 0 REU) defined which mutations will be included in each combinatorial design based on the previous energy calculations. The PROSS algorithm was run with default parameters using open or closed state crystal structures of each PBP as input. Generally, the binding pocket residues 6 Å around the ligands in the closed structures were held fixed in all designs to ensure that interactions with the ligands are kept intact. In the case of Design X.4, hinge point and interface residues were fixed (see Methods below - Gaussian network models). Designs with the same Rosetta energy cut-off and 4-10 % mutational load were chosen. Each mutation was inspected and hand-picked mutations were excluded if mutations were too close to the termini or no rationale of stabilizing the protein could be found. If the ΔΔ*G* of a mutation was stabilizing (ΔΔ*G* < 0) in both the open and closed state designs, this mutation was included in the two-state design variants (two-state filter). Categorization of mutations was done with the Rosetta LayerSelector using default settings.

### Gaussian network models

GNMs were previously used to derive models of protein backbone flexibility based on Cα atom positions that correlated with crystallographic B-factors but are not restricted to experimentally determined structures.^43–47^ PDBs of the open and closed ligand-bound structure were used (Table S1). Ligands were deleted and in the case of TphC, the residue numbering of the closed state was adjusted to that of the open state. The per-residue square fluctuations were obtained by GNM calculations with the distance cut-off set to 10 Å and the spring constant set to 1 using the ProDy python implementation.^48,49^ The number of modes was increased until no further features were observed. 5 modes was the lowest number that reflected the description of PBP dynamics.

To obtain the hinge residues, GNM was calculated based on only the 1st mode. The ProDy python implementation provides a function to output hinge residues. The protein lobes were then split along the hinge residues and the interface residues between the lobes were determined with a 6 Å distance cut-off using the PyMol^50^ python implementation.

### Cloning, expression & purification

PBPs are naturally expressed with a signal sequence for the transport to the periplasm of gram-negative bacteria. As a previously reported protein purification protocol was used for all PBPs tested here ^28^, the signal sequences were removed thus leading to an expression in the cytoplasm. The codon-optimized genes were synthesized by Twist Bioscience HQ including restriction sites for *Nde*I and *Xho*I on the N-terminal and C-terminal flanking sites, respectively (see Table S2 for amino acid sequences). The DNA fragments were first digested and then ligated into a pET21b(+)-vector which included a C-terminal His-tag. In case of TphC fragments, a N-terminal His-tag and TEV-cleavage site was included in the synthesized gene because a previous study demonstrated successful expression and purification with the His-tag at this position.^35^ A stop codon was inserted between the gene and the *Xho*I cleavage site by PCR (primers listed in Table S3) before cloning into the expression vector. Top10 *E. coli* cells were transformed with the generated vectors and resulting clones were verified by Sanger sequencing. For expression, BL21 (DE3) cells were transformed with the plasmids and plated out on LB agar plates containing 100 µg/mL ampicillin. Each 25 mL LB media-based overnight culture supplemented with 100 µg/mL ampicillin was inoculated with a single colony. 0.5 L TB media was then inoculated with 5 mL of the overnight culture and incubated at 37 ℃ until the OD_600_ reached a value of ∼ 0.7. Next, overexpression was induced by adding isopropyl-ꞵ-thiogalactoside (IPTG) to a final concentration of 1 mM and cultures were incubated for ∼ 18 h at 18 ℃. Cells were harvested by centrifugation (Beckman Coulter, Avanti JXN-26, JLA8.1000 rotor) at 4,000 g, 4 ℃ for 20 min. The cell pellets were washed with 30 mL lysis buffer (25 mM NaH_2_PO_4_/Na_2_HPO_4_, 500 mM NaCl, 20 mM Imidazole, pH 7.4) supplemented with protease inhibitor mix HP (Serva) and centrifuged again (Eppendorf 5920R) at 4,000 g, 4 ℃ for 10 min. The cell pellets were resuspended in lysis buffer, sonicated (Branson 6.3 mm tip, 2 x 3 min, 40 % duty cycle, output power 4) and centrifuged (Beckman Coulter, Avanti JXN-26, JA25.50 rotor) at 18,000 g, 4 ℃ for 1 h. The supernatant was loaded onto a His-Trap HP 5 mL column (GE Healthcare) that was equilibrated with the lysis buffer beforehand using a peristaltic pump (Pump P-1, Pharmacia). After loading, the column was washed with 10 column volumes (CV) of lysis buffer. Next, the protein was unfolded by adding 5 CV 6 M Guanidine Hydrochloride (GdHCl) and incubation for 1 h. Endogenous ligands were removed by washing with 5 CV GdHCl. The protein was refolded by washing with 10 CV of lysis buffer. Elution was performed in a stepwise manner on an ÄKTA system using elution buffer (25 mM NaH_2_PO_4_/Na_2_HPO_4_, 500 mM NaCl, 500 mM Imidazole, pH 7.4). The protein fractions were pooled and concentrated using centrifugal concentrators to a final volume of 10 mL and loaded onto a preparative size exclusion column (HiLoad Superdex 75 26/60, GE Healthcare) equilibrated with running buffer (25 mM NaH_2_PO_4_/Na_2_HPO_4_, 500 mM NaCl, pH 7.4). Monomeric protein fractions were pooled and concentrated for the subsequent measurements. Concentrations were checked using a NanoDrop Spectrophotometer (Eppendorf BioPhotometer D30). Expressions and purifications were validated by SDS-PAGE.

### Circular dichroism spectroscopy

To remove salts, protein samples were dialyzed overnight against 2 L of CD buffer (25 mM NaH_2_PO_4_/Na_2_HPO_4_, pH 7.4) using a 10 kDa molecular weight cut-off. Aggregated protein was removed by centrifugation at 15,000 g for 5 min using a tabletop centrifuge (Eppendorf 5427R). Next, the protein concentration was adjusted to 0.3 mg/mL and verified photometrically. 200 µL of protein samples were mixed with either 100 µL CD buffer or 100 µL of ligand solution (300 µM stocks of ligand solubilized in CD buffer) to reach a final concentration of 100 µM for the ligands and 0.2 mg/ml for the protein. CD spectra and thermal melt experiments were obtained on a JASCO J-1500 using a glass cuvette with a 1 mm path length. For CD spectra, following settings were used: wavelengths 240-190 nm, bandwidth 1 nm, response time 2 s, data pitch 0.1 nm, scanning speed 100 nm/min and ten accumulations (for buffer controls only one accumulation was used). For Tmelt measurements, the following settings were used: wavelength of 222 nm, temperature range from 20-95 ℃ controlled by a Pelletier AWC100 (Julabo) using a slope of 1 ℃/min, a bandwidth of 1 nm and a response time of 2 s. Raw CD data was normalized by subtraction of the respective buffer data and subsequent conversion of the measured signal in millidegree to mean residue molar ellipticity.^51^ To obtain the apparent melting temperature *T_m_*, thermal melt data was fit using a sigmoidal equation according to Niklasson et al.^52^ The error plotted stems from the fit function uncertainty.

### Isothermal titration calorimetry

Ligands were dissolved in the specific buffer that was used in the size-exclusion chromatography (SEC) run of each protein purification (stored at - 20 ℃ and thawed on the day of use). All protein and ligand samples were degassed and temperature equilibrated to 25 ℃ for at least 10 min using a degassing station (TA Instruments). 400 µL of protein samples were loaded into the sample cell of a low volume AffinityITC machine with gold cell (TA Instruments) and ligand solutions were added into the injection syringe (protein and ligand concentrations can be found in Table S5). ITC measurements were performed at 25 ℃ with 20 injections of 2.5 µL (one initial injection of 0.5 µL was excluded from analysis), an adaptive injection interval and stirring rate of 125 rpm. Subtractions of constant heat of dilution values for each ligand, peak integration and fitting with a one-site binding model were done with NanoAnalyze (TA Instruments). The reported errors reflect the standard deviation of three technical replicas. Thermograms and binding isotherms were plotted in Figures S10-13.

## Author Contributions

Conceptualization – M.S., B.H. & S.J.F., Investigation – M.S., Formal Analysis – M.S. & M.K., Manuscript writing – all authors.

## Acknowledgments

We thank Ariel Tennenhouse for adding suggestions on the manuscript. We thank Sabrina Wischt and Pascal Kröger for experimental support. This project has received funding from the European Union’s Horizon 2020 research and innovation programme under the Marie Skłodowska-Curie grant agreement No. 955623 (to B.H. and S.J.F.) and the European Synergy grant agreement No 951375 (ArtMotor) (to B.H.).

## Abbreviations

PBP: periplasmic binding protein
PotF: putrescine binding protein
TphC: terephthalate binding protein
MBP: maltose binding protein
LAO: L-lysine/L-arginine/L-ornithine binding protein

## Supporting Information

### Supporting Tables

**Supporting Table 1.**
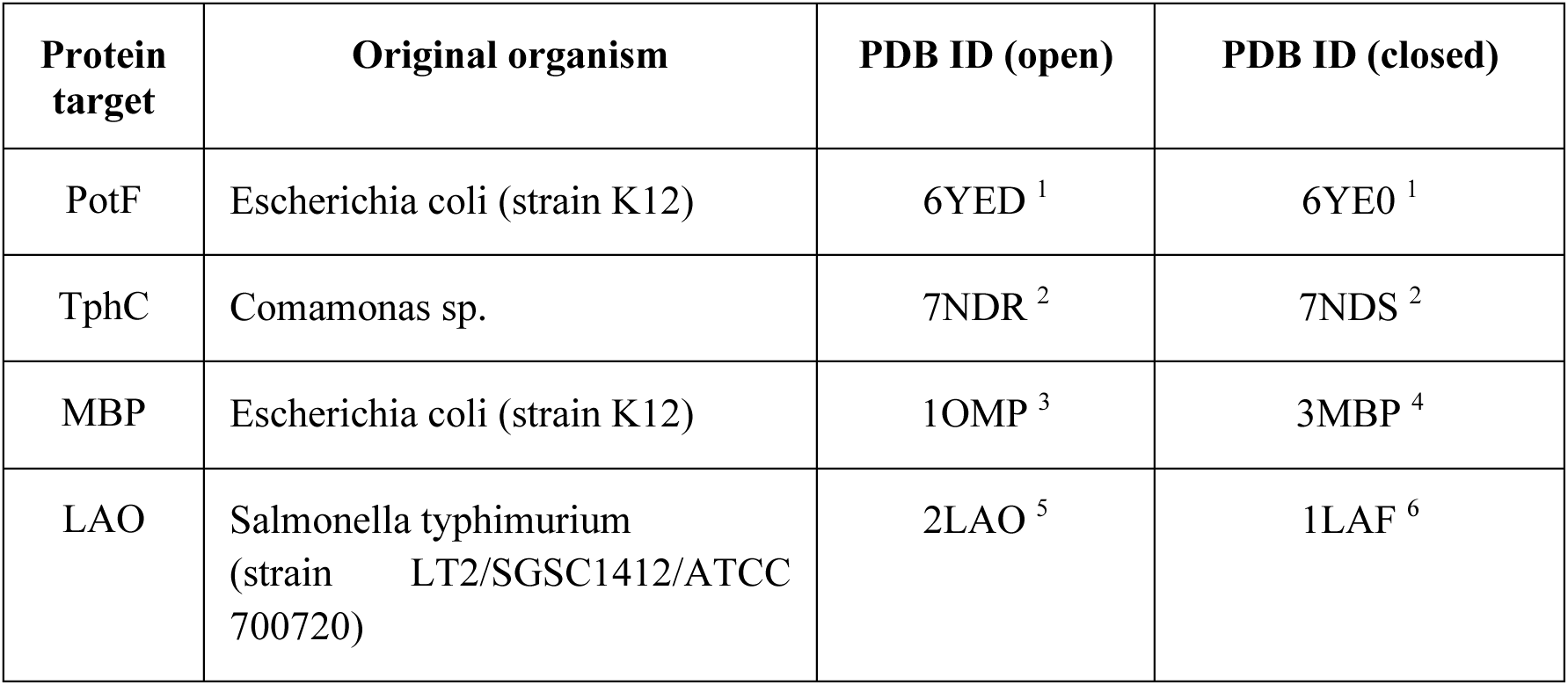
Original organism and PDB^7^ identifiers of the PBP crystal structures used.

**Supporting Table 2.**
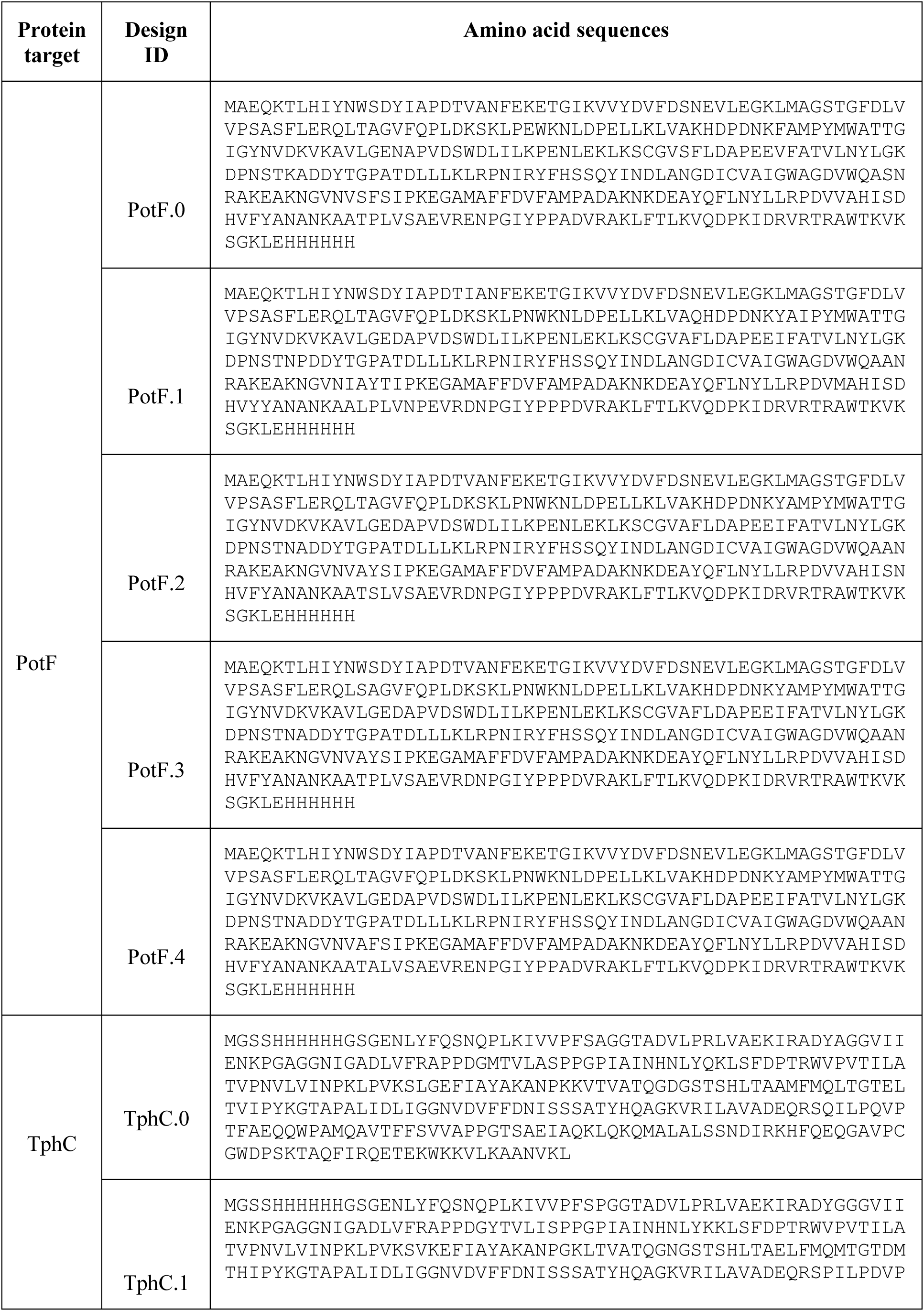
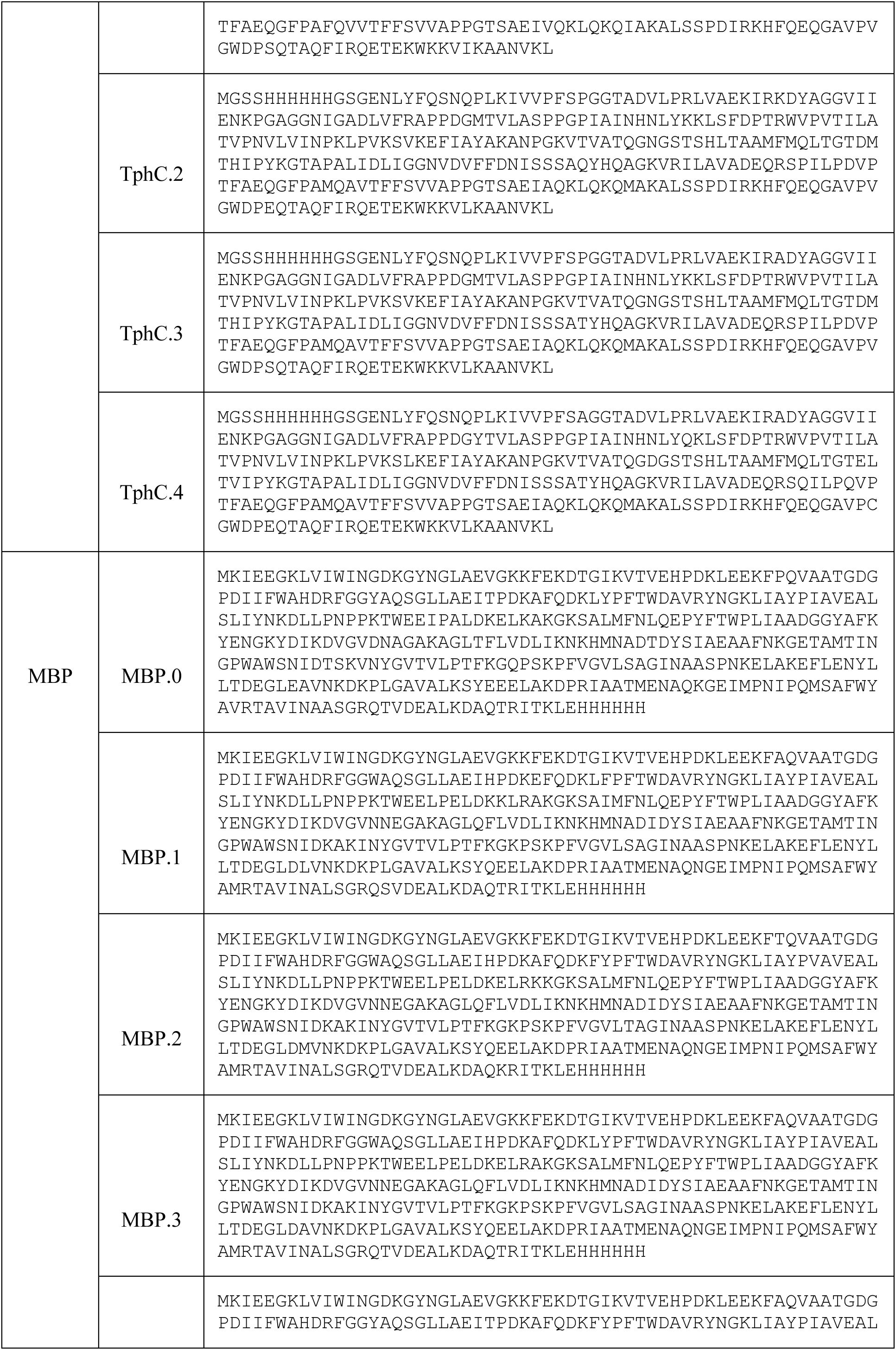
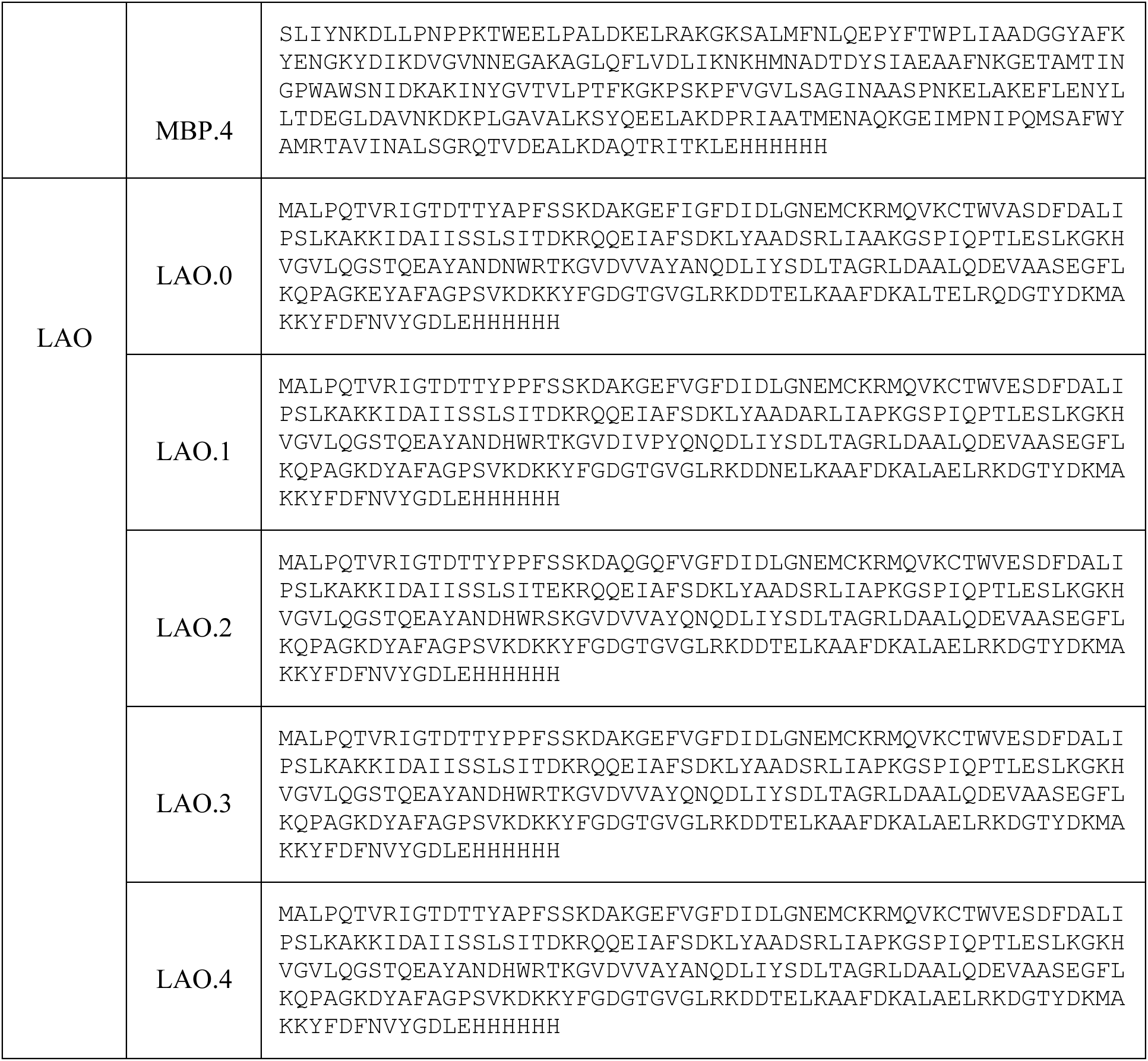
Amino acid sequences of PotF, TphC, MBP and LAO WT and designs tested in this study.

**Supporting Table 3.**
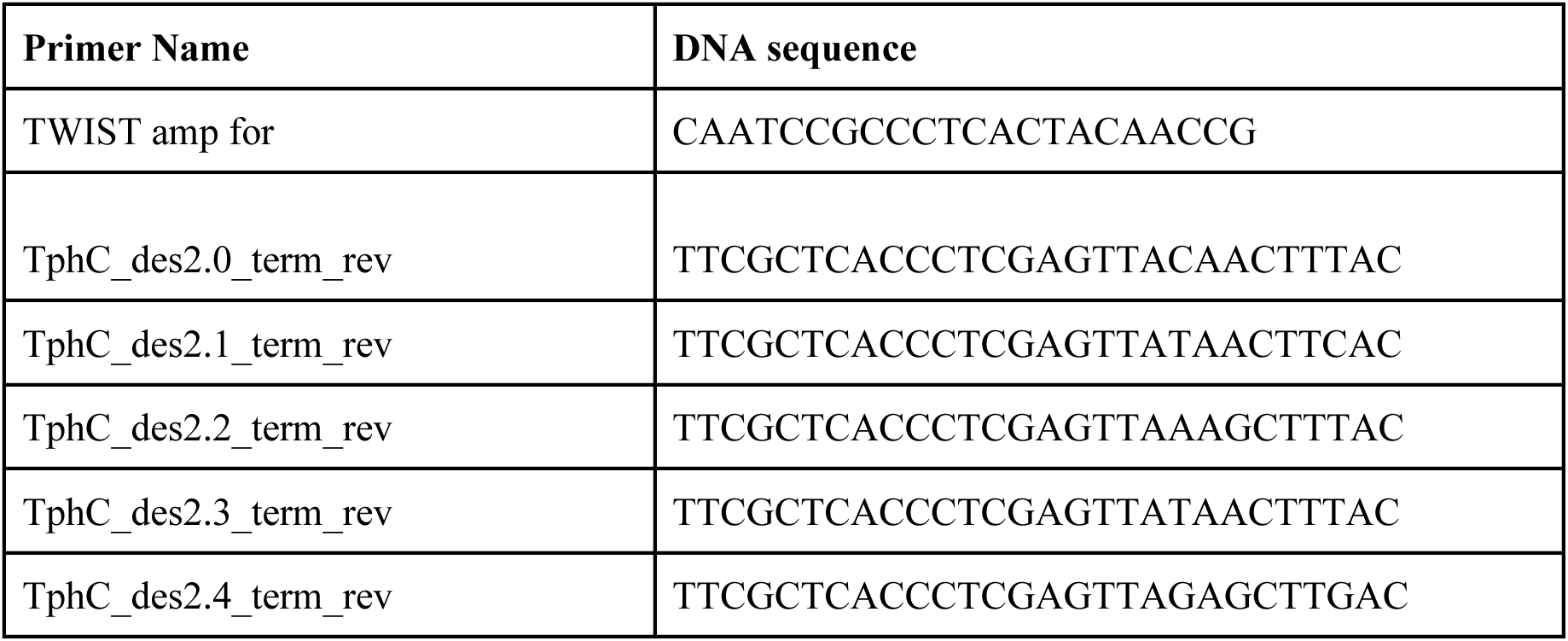
Primers used in this study.

**Supporting Table 4.**
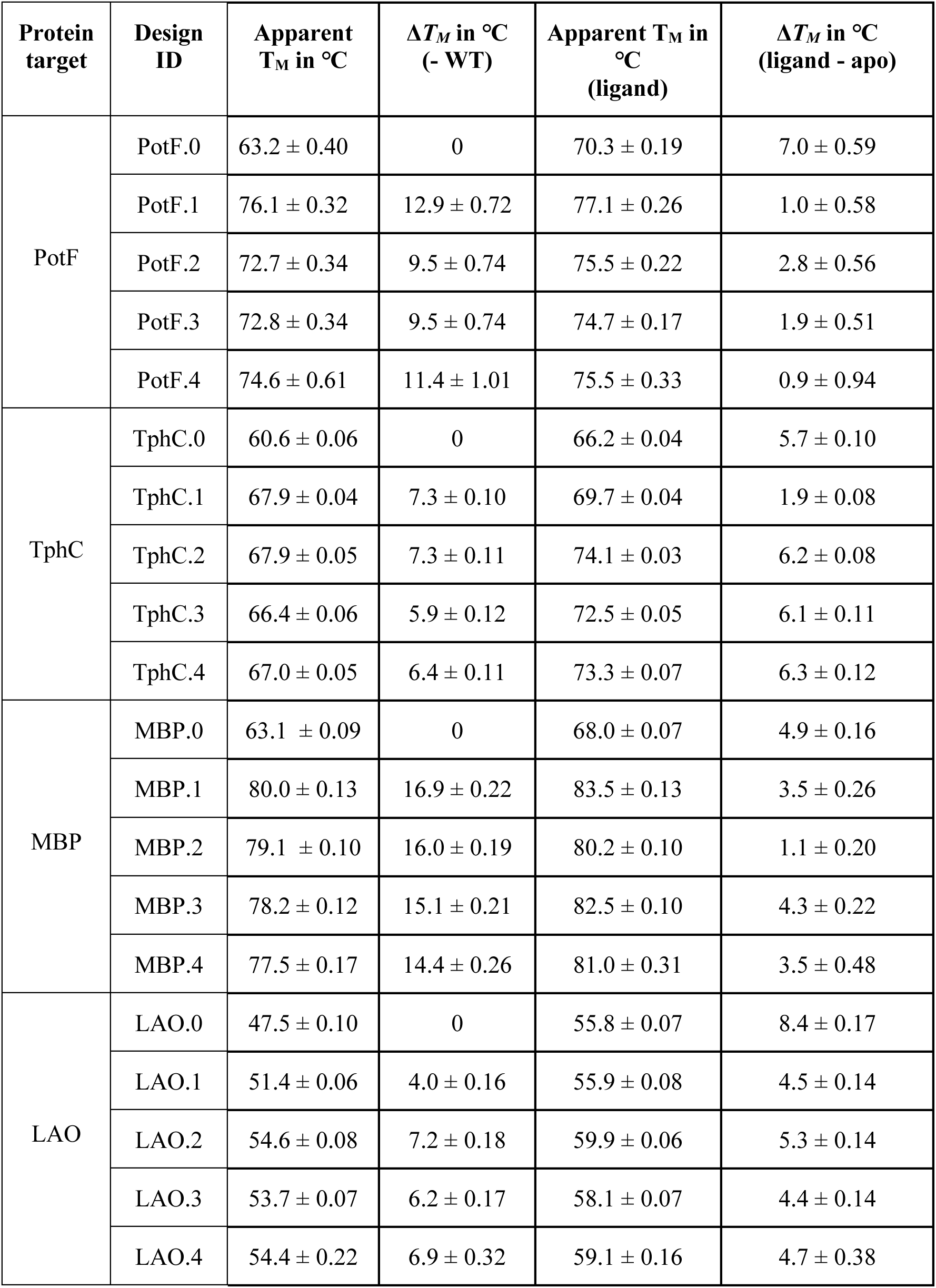
Thermal melt data of all PBP variants.

**Supporting Table 5.**
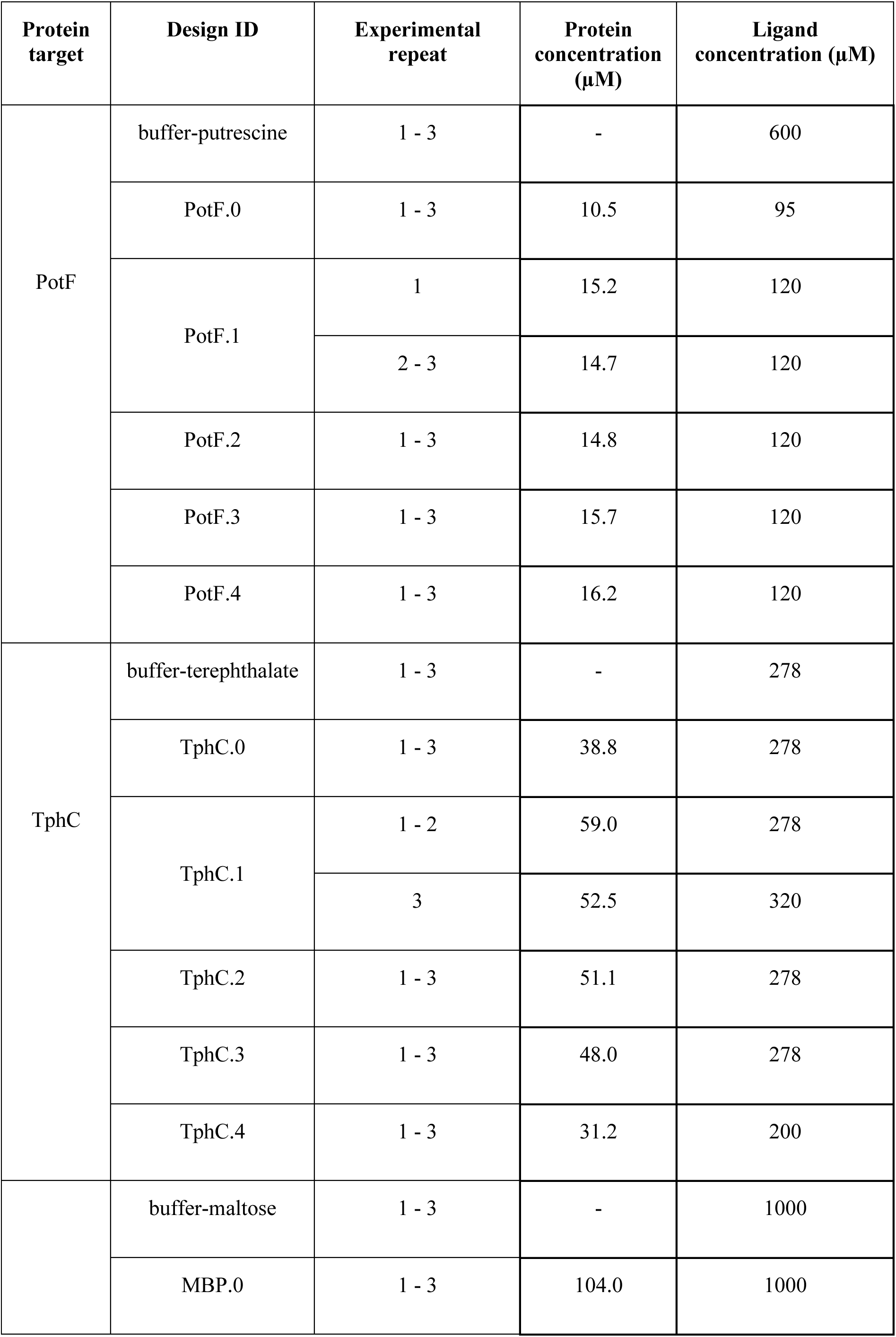
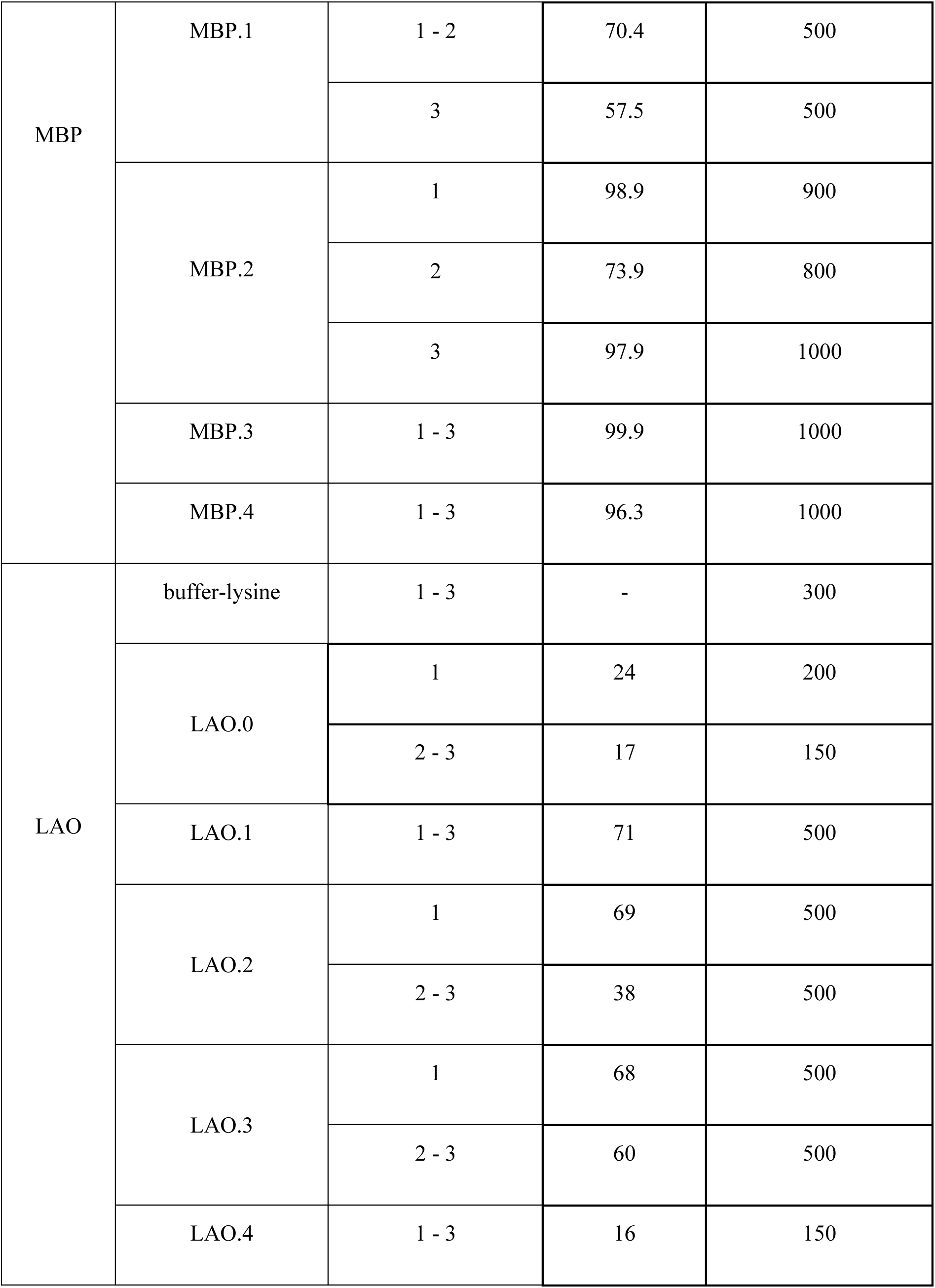
Protein and ligand concentrations for ITC measurements.

**Supporting Table 6.**
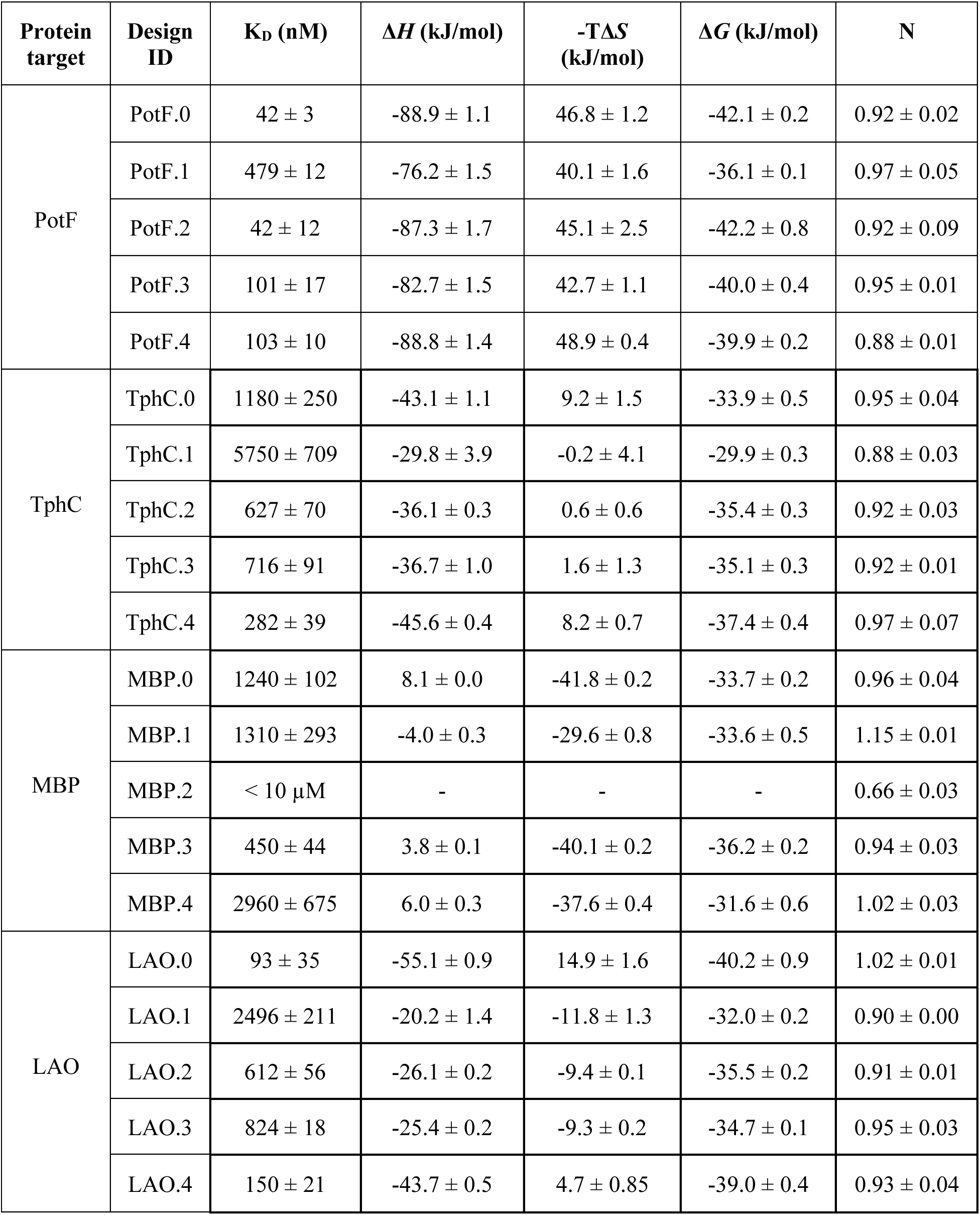
ITC data of all PBP variants. Averages and standard deviations were calculated based on three technical replicas.

### Supporting Figures

**Supporting Figure 1.**
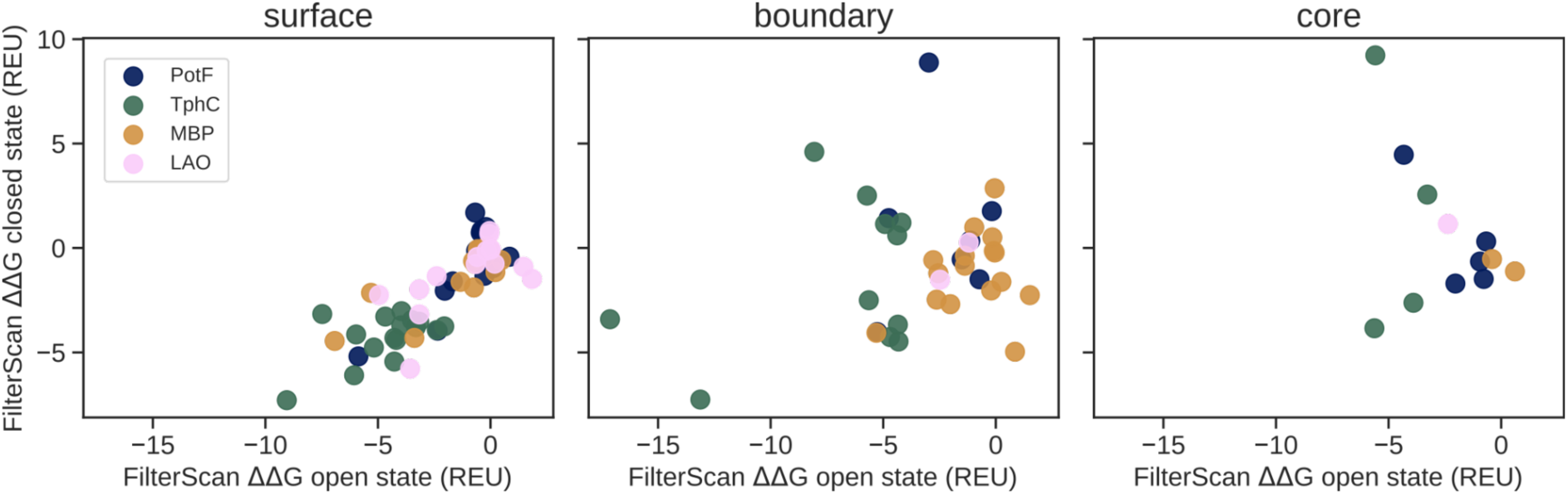
ΔΔ*G* values of PROSS mutations generated of PotF, TphC, MBP and LAO filtered by Rosetta LayerSelector categories: surface, boundary, and core. Open- and closed-state energies were plotted against each other.

**Supporting Figure 2.**
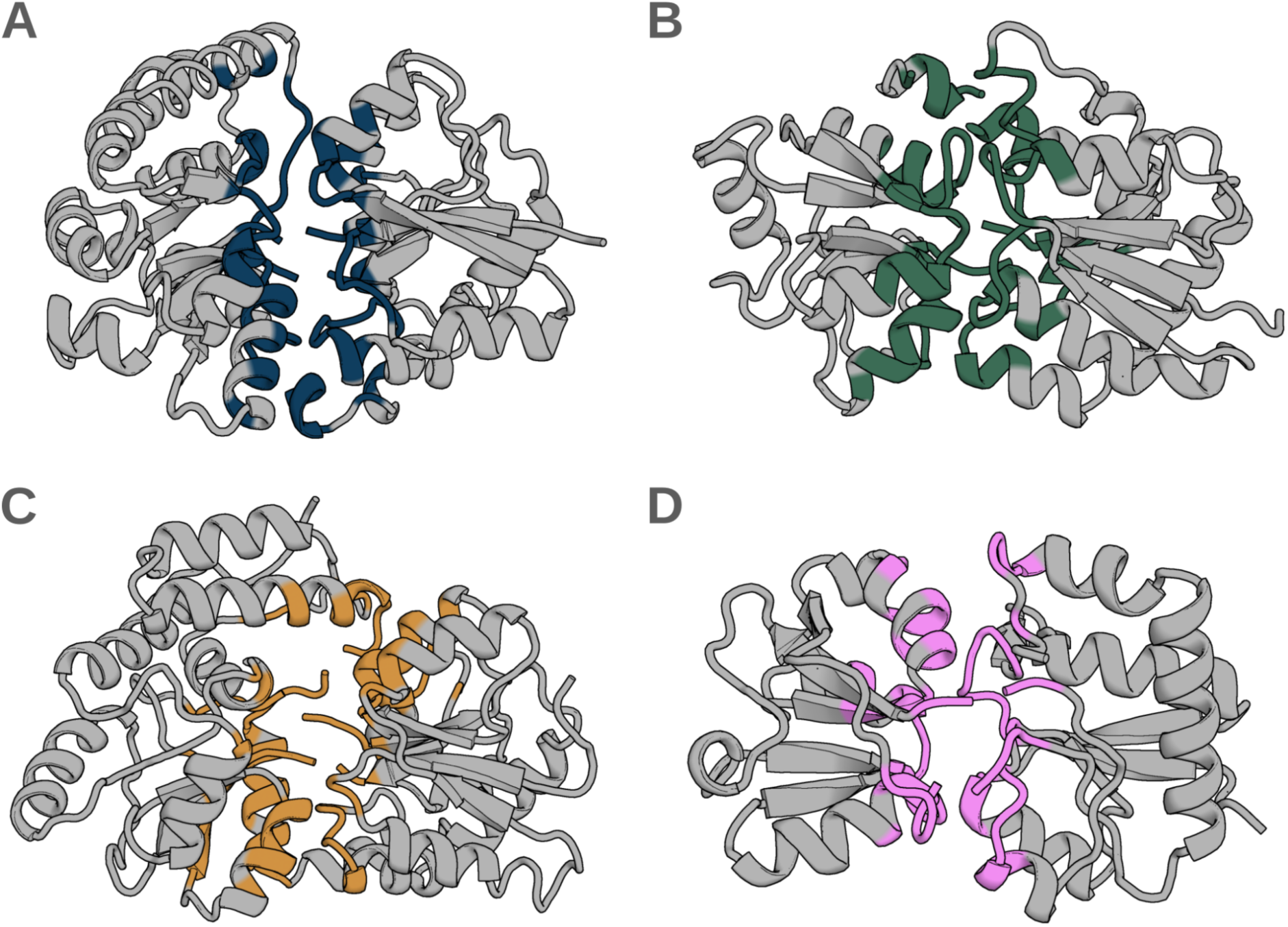
PBP interface residues that were fixed during PROSS design. (A-D) Closed-state PotF, TphC, MBP and LAO structures were split by predicted hinge residues and subdomain interface residues were highlighted in blue, green, orange and pink, respectively.

**Supporting Figure 3.**
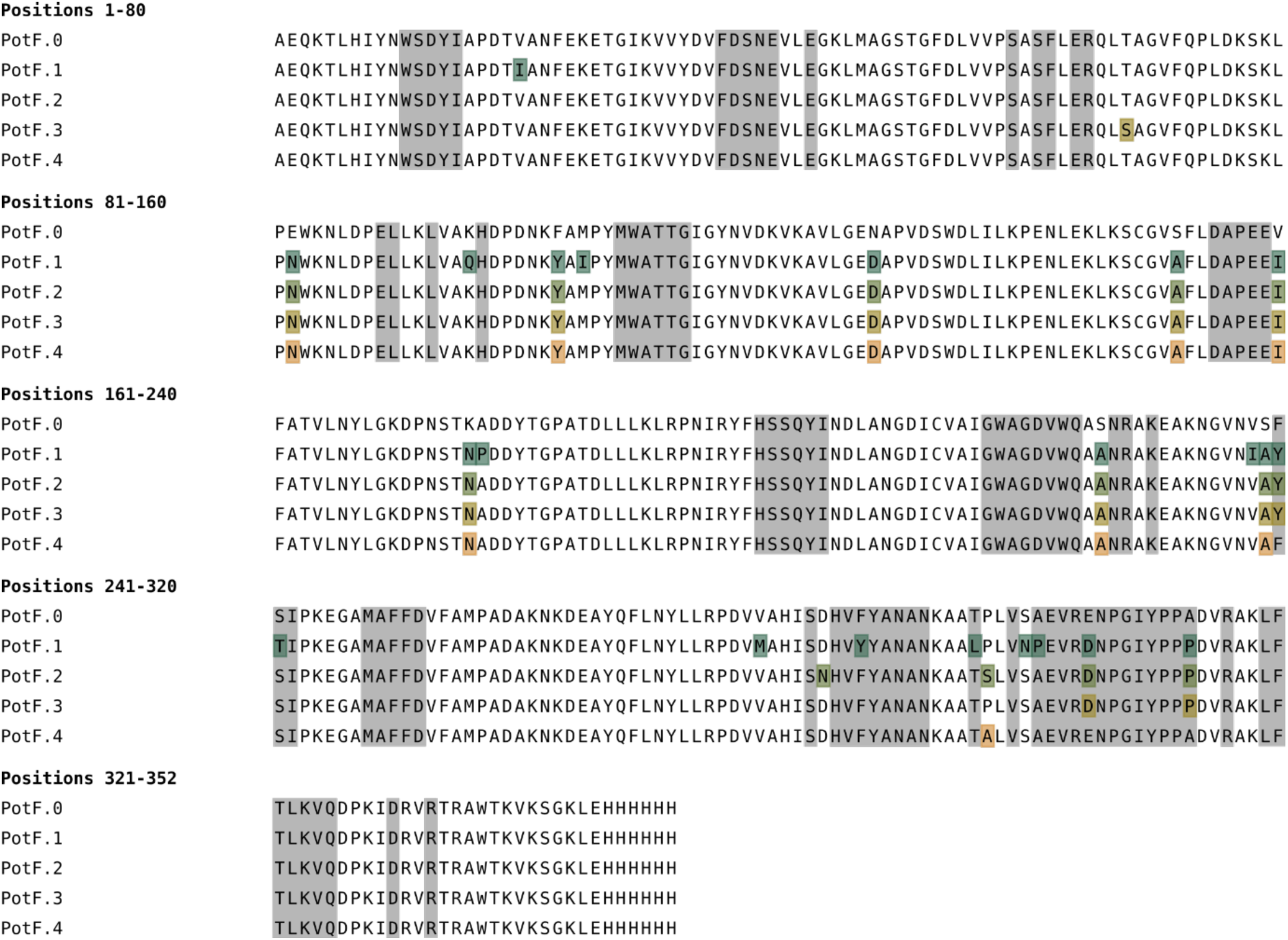
Amino acid sequence alignment of PotF.0-PotF.4 with mutations highlighted. Additionally, hinge and interface residues are shown as grey boxes.

**Supporting Figure 4.**
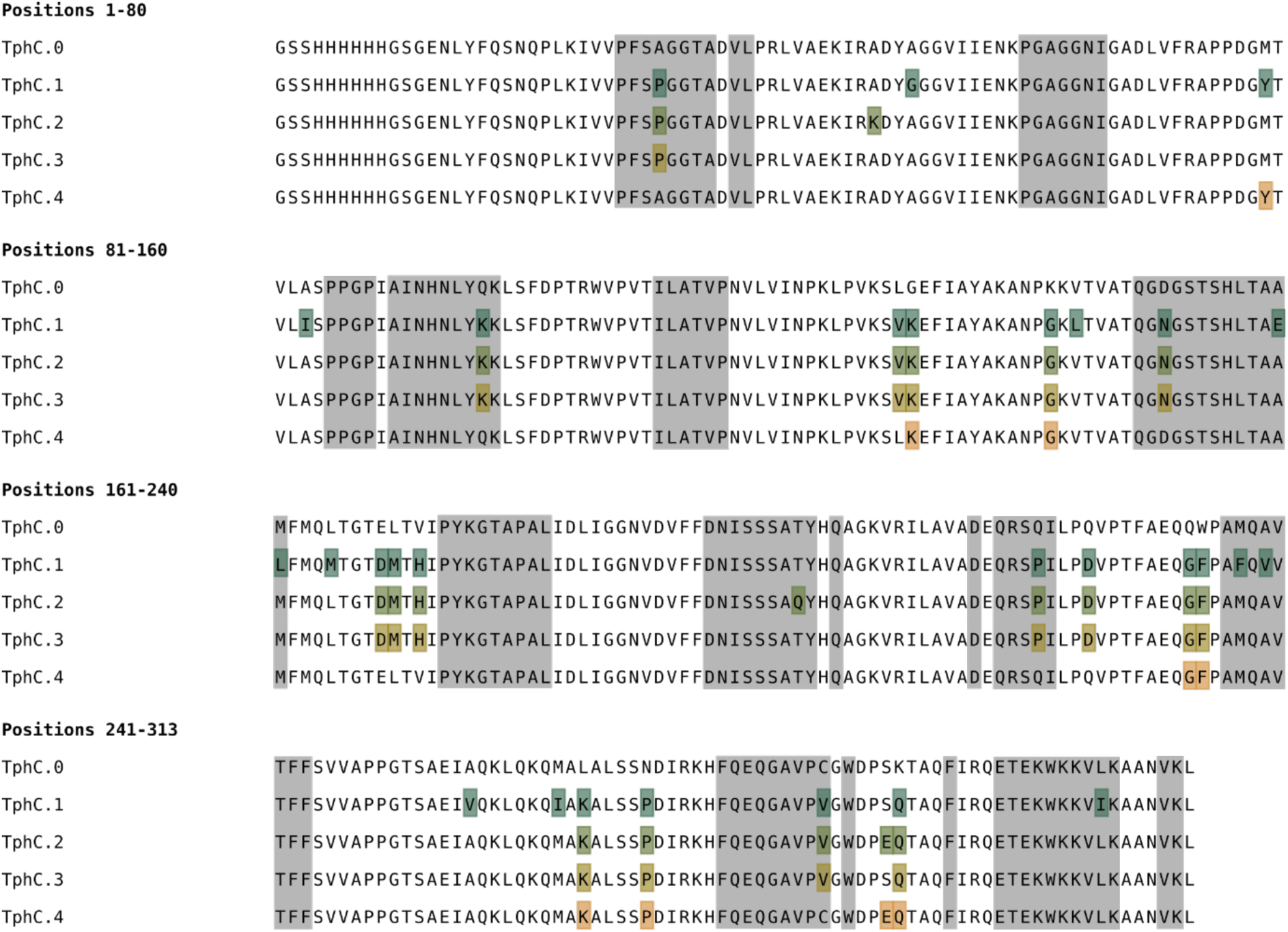
Amino acid sequence alignment of TphC.0-TphC.4 with mutations highlighted. Additionally, hinge and interface residues are shown as grey boxes.

**Supporting Figure 5.**
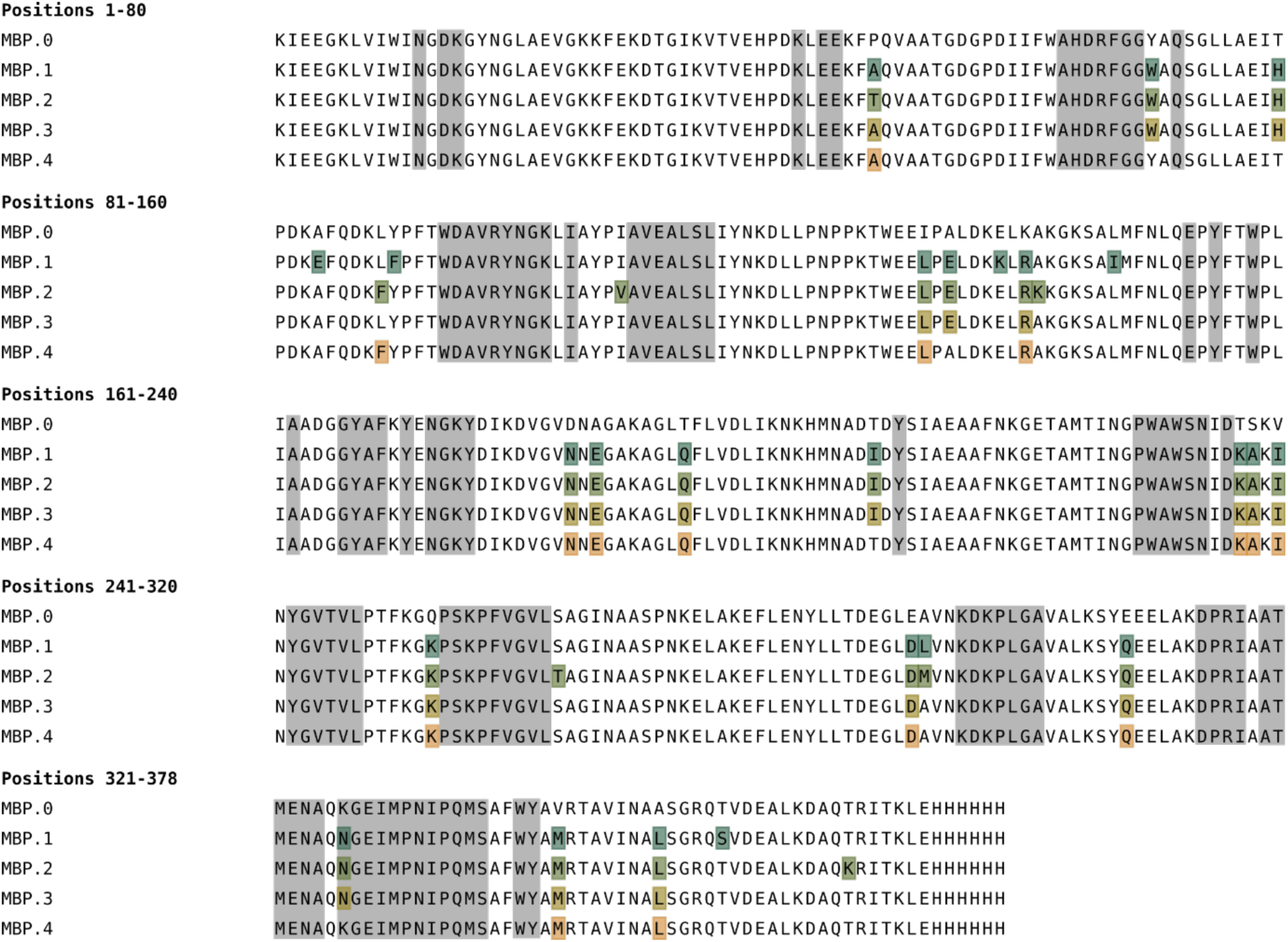
Amino acid sequence alignment of MBP.0-MBP.4 with mutations highlighted. Additionally, hinge and interface residues are shown as grey boxes.

**Supporting Figure 6.**
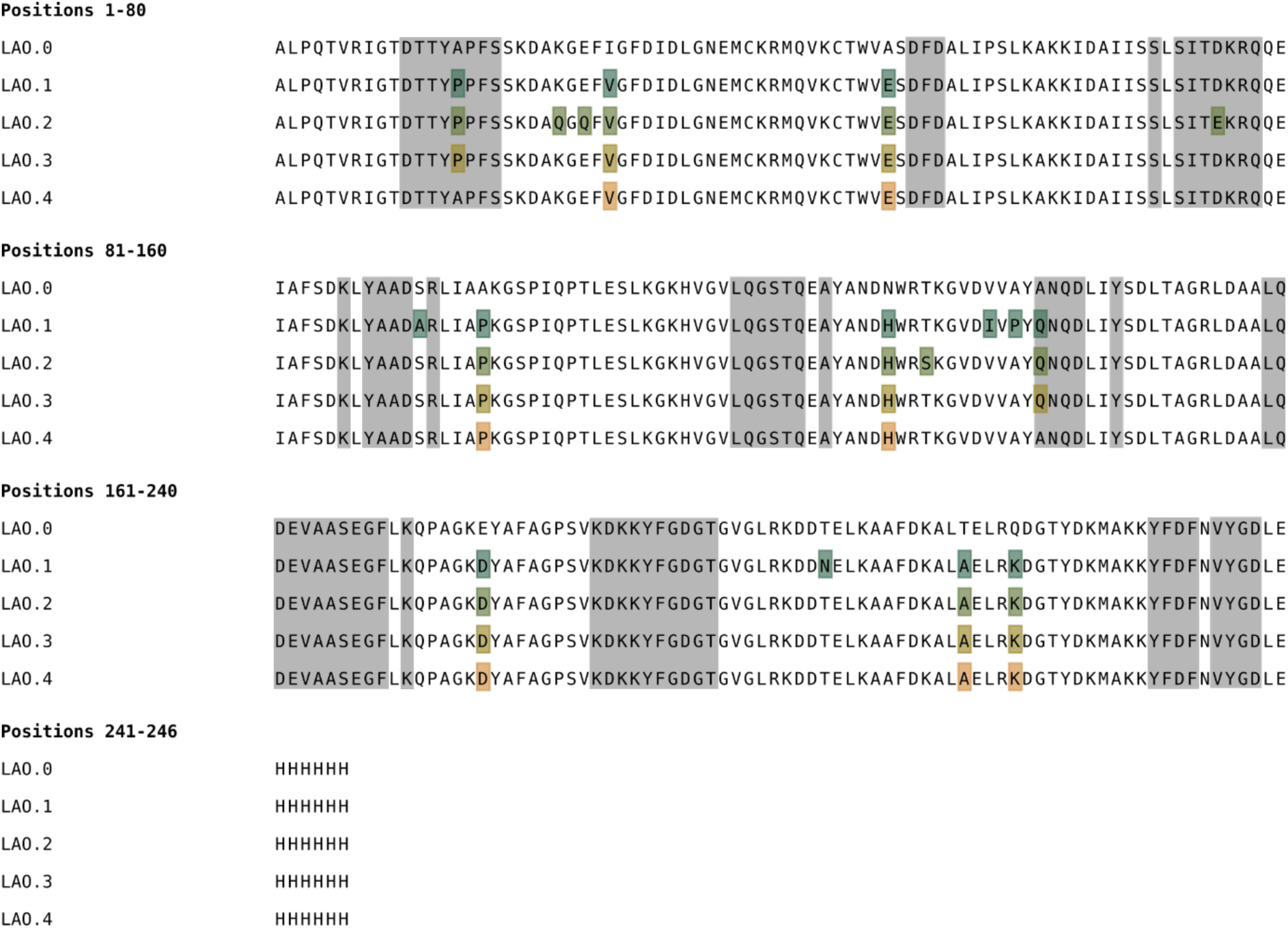
Amino acid sequence alignment of LAO.0-LAO.4 with mutations highlighted. Additionally, hinge and interface residues are shown as grey boxes.

**Supporting Figure 7.**
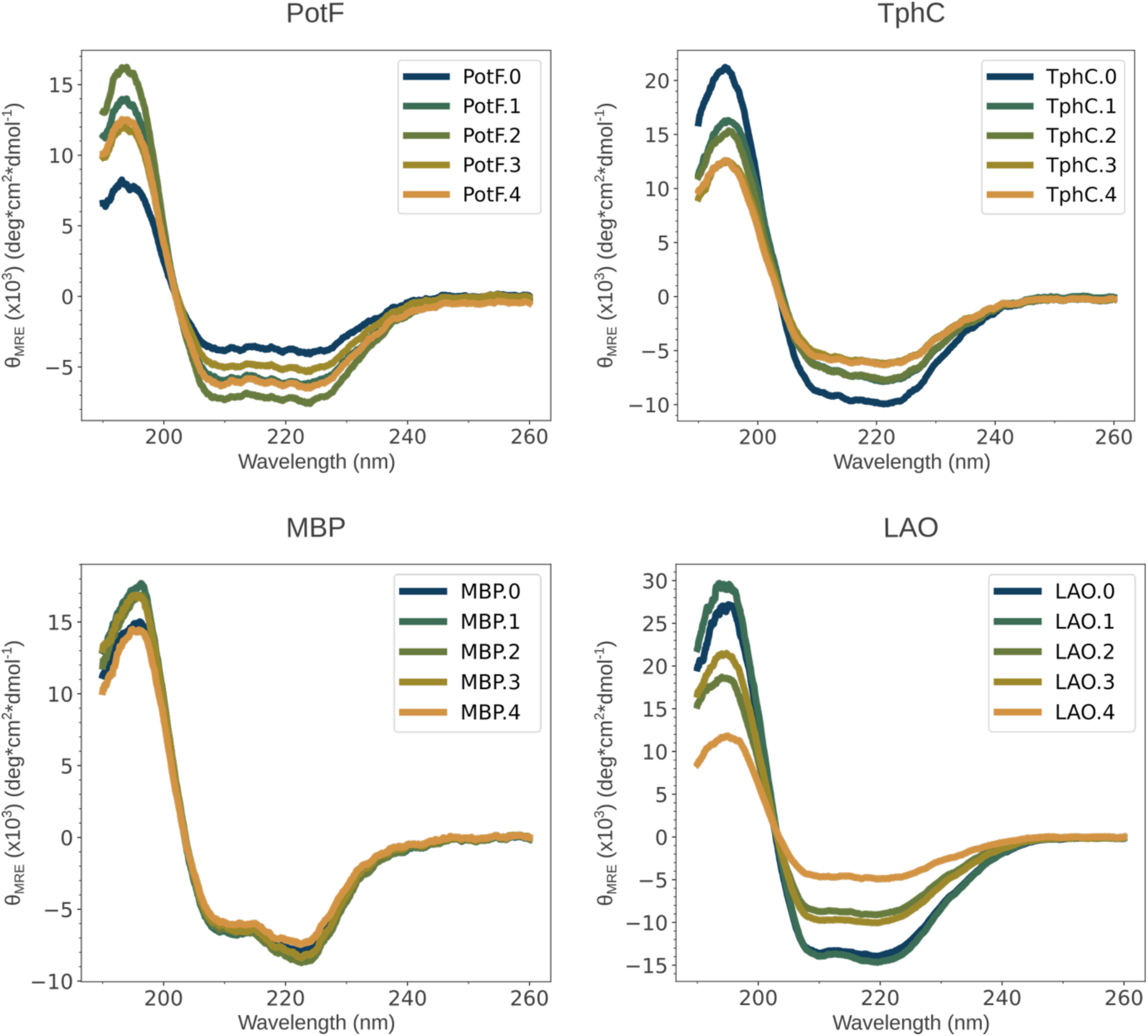
CD spectra of all PBP variants without ligands present plotted as mean residue ellipticity (θ_MRE_) against the wavelength.

**Supporting Figure 8.**
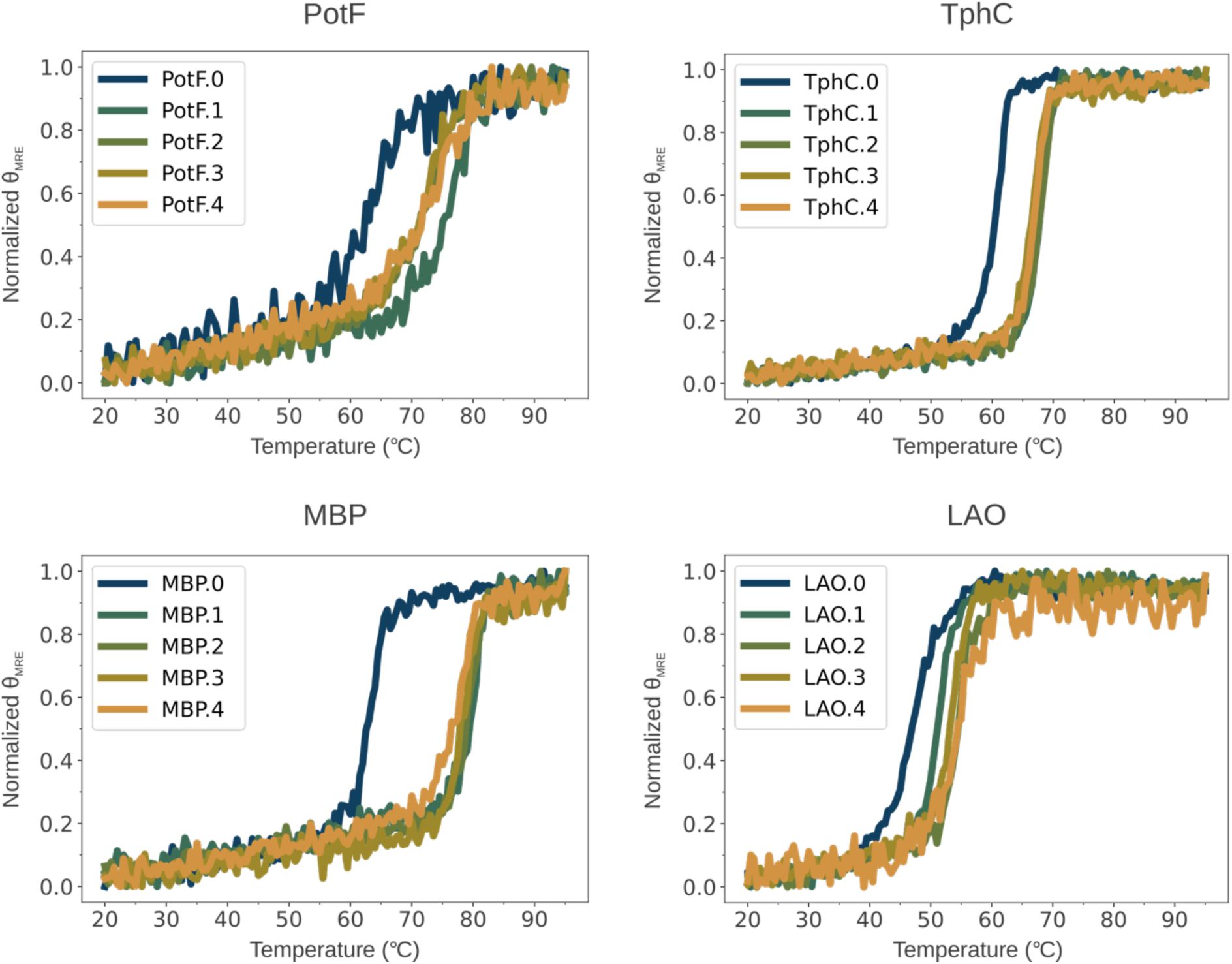
CD thermal melts of all PBP variants without ligands present plotted as normalized mean residue ellipticity (θ_MRE_) against a temperature gradient.

**Supporting Figure 9.**
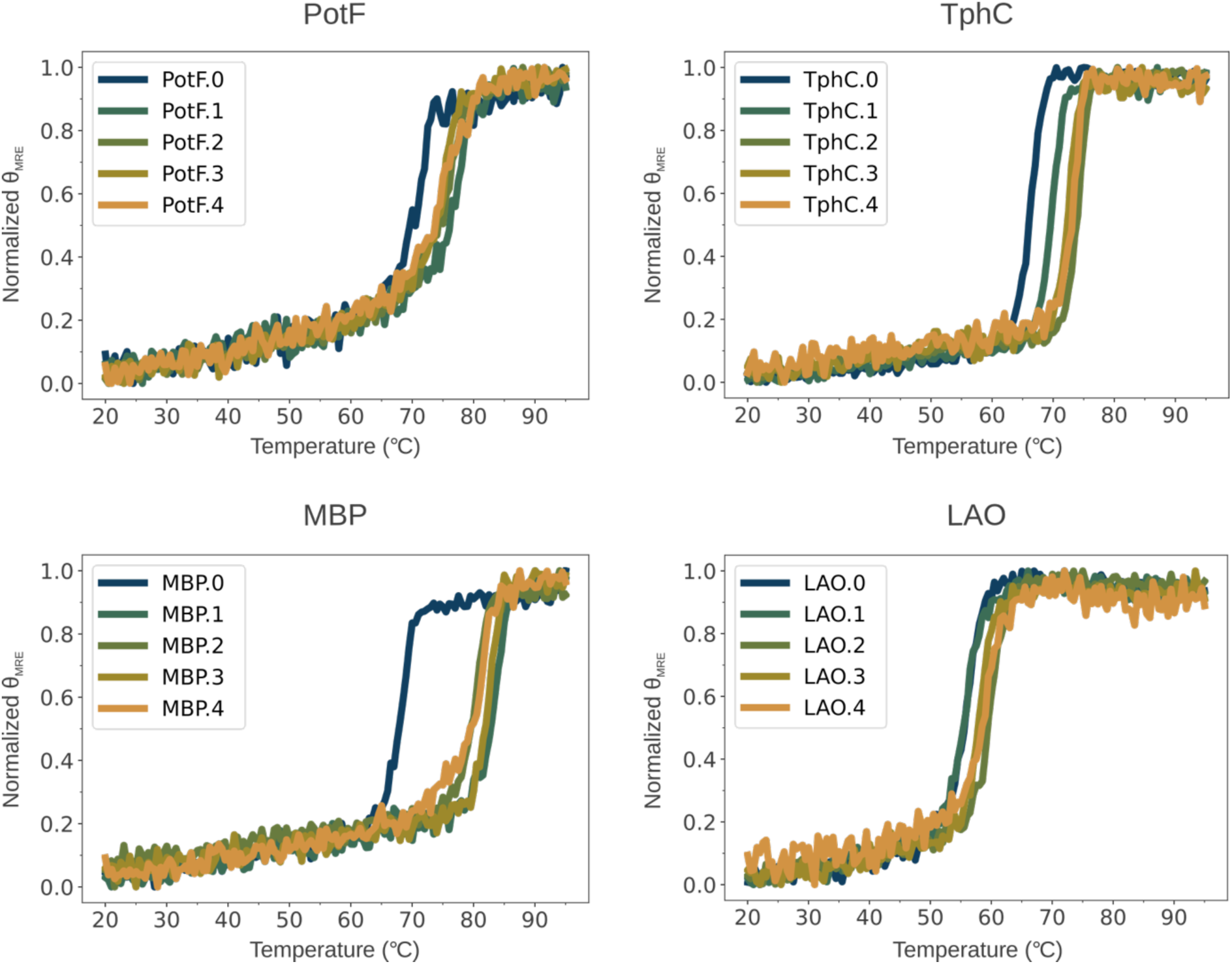
CD thermal melts of all PBP variants with ligands present plotted as normalized mean residue ellipticity (θ_MRE_) against a temperature gradient.

**Supporting Figure 10.**
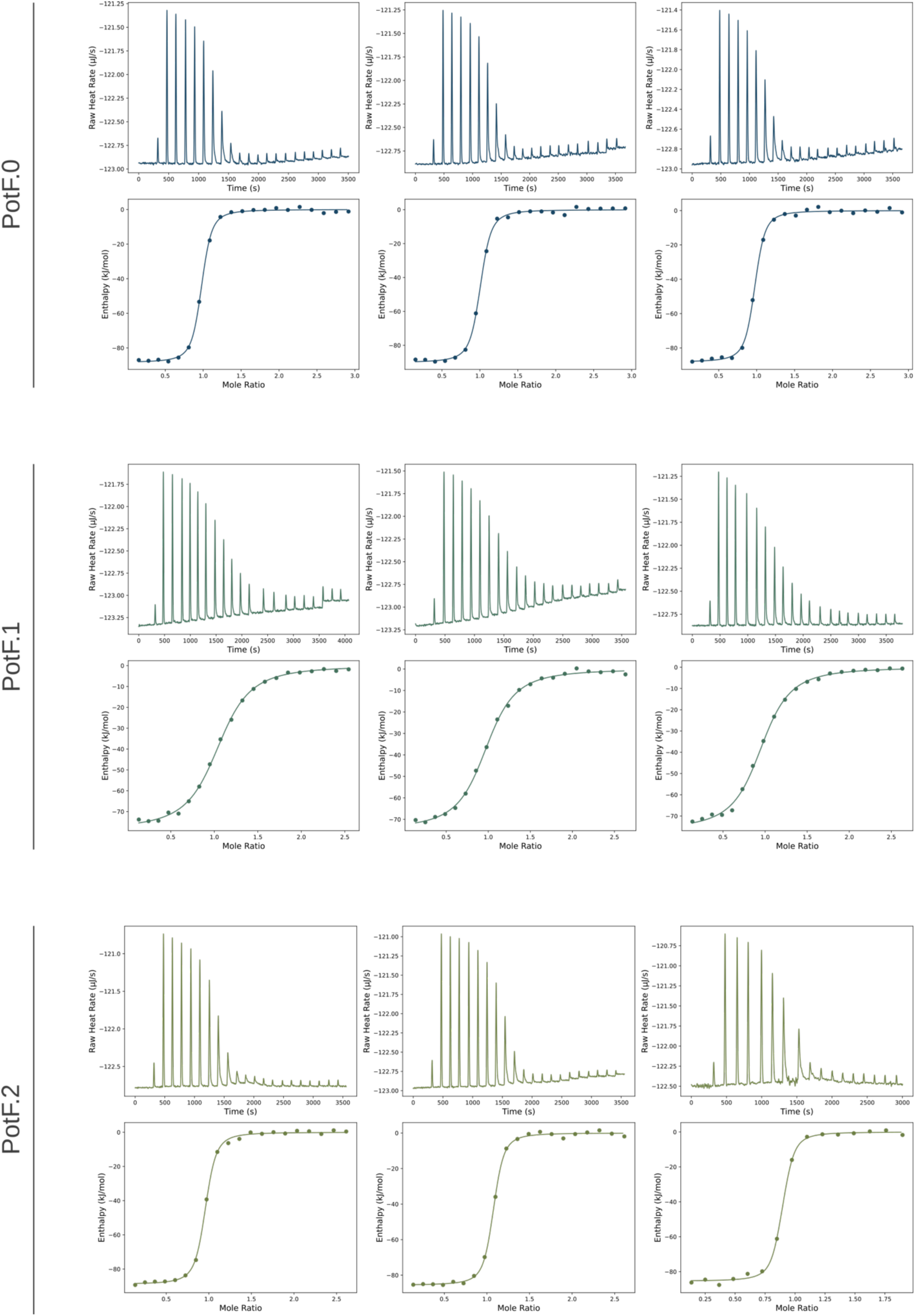
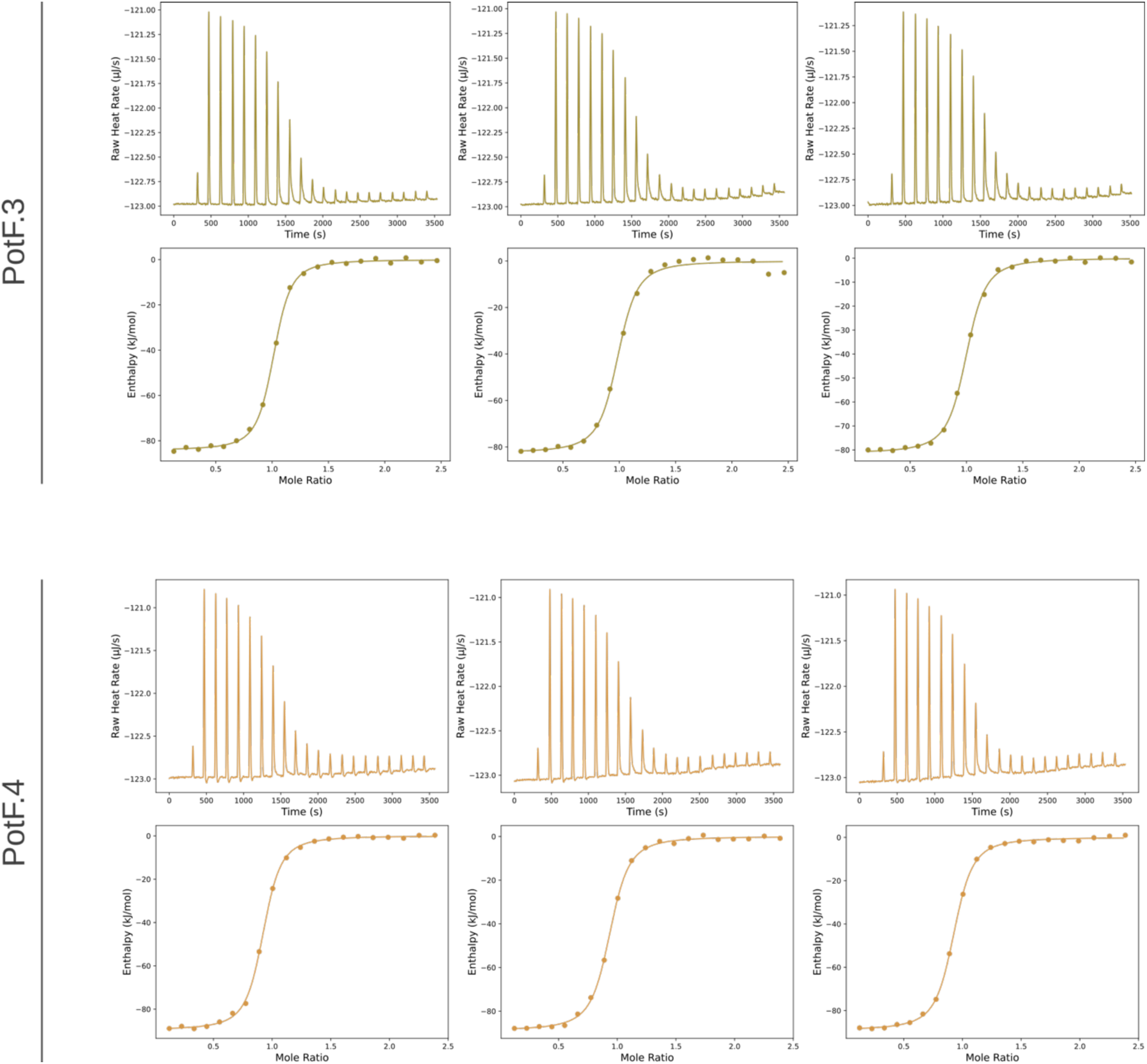
Thermograms and binding isotherms of PotF.0-4 and putrescine in three technical replicates. Buffer-putrescine control titrations were performed to obtain baseline values that were subtracted from the integrated heats of PotF.0-4.

**Supporting Figure 11.**
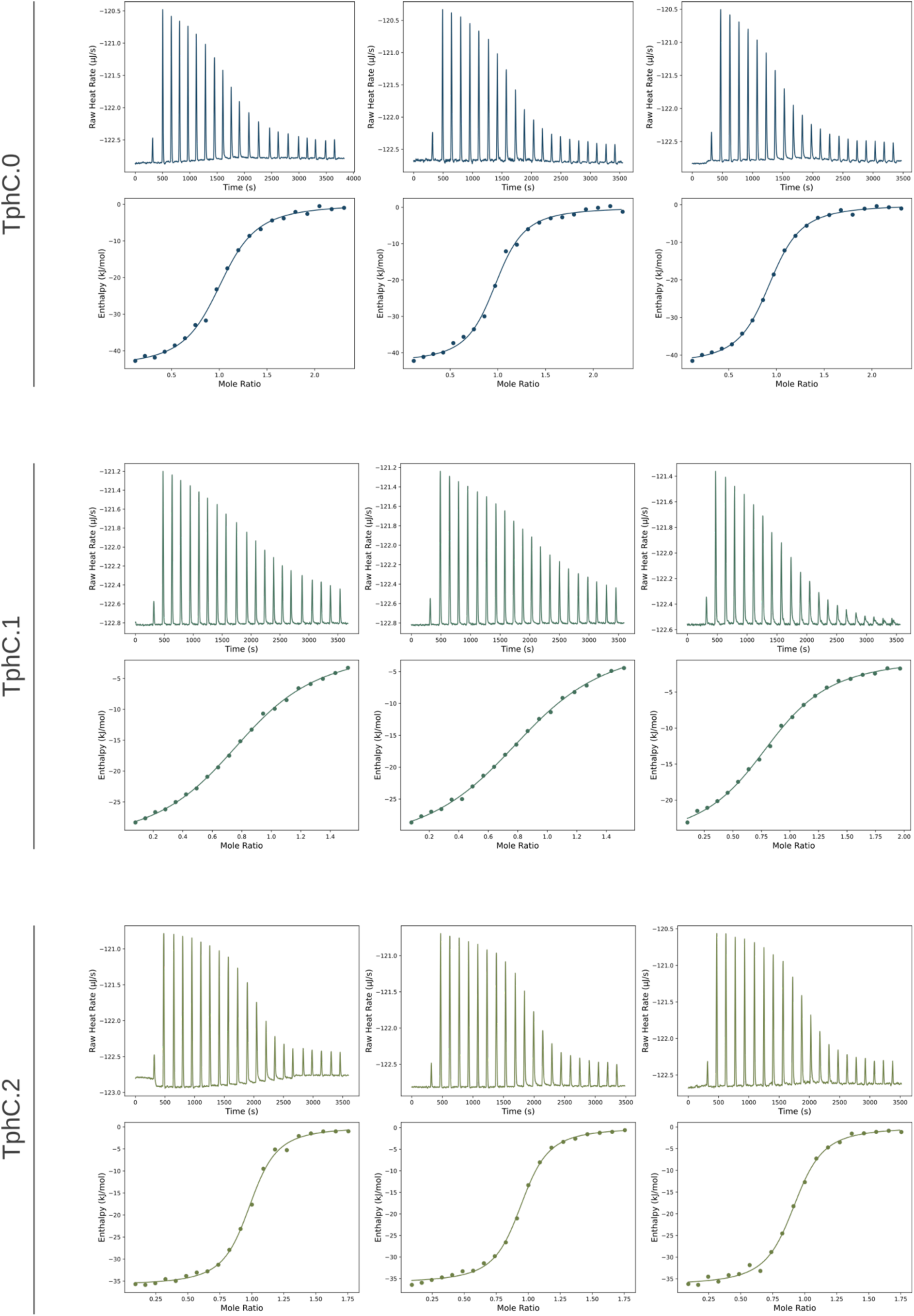
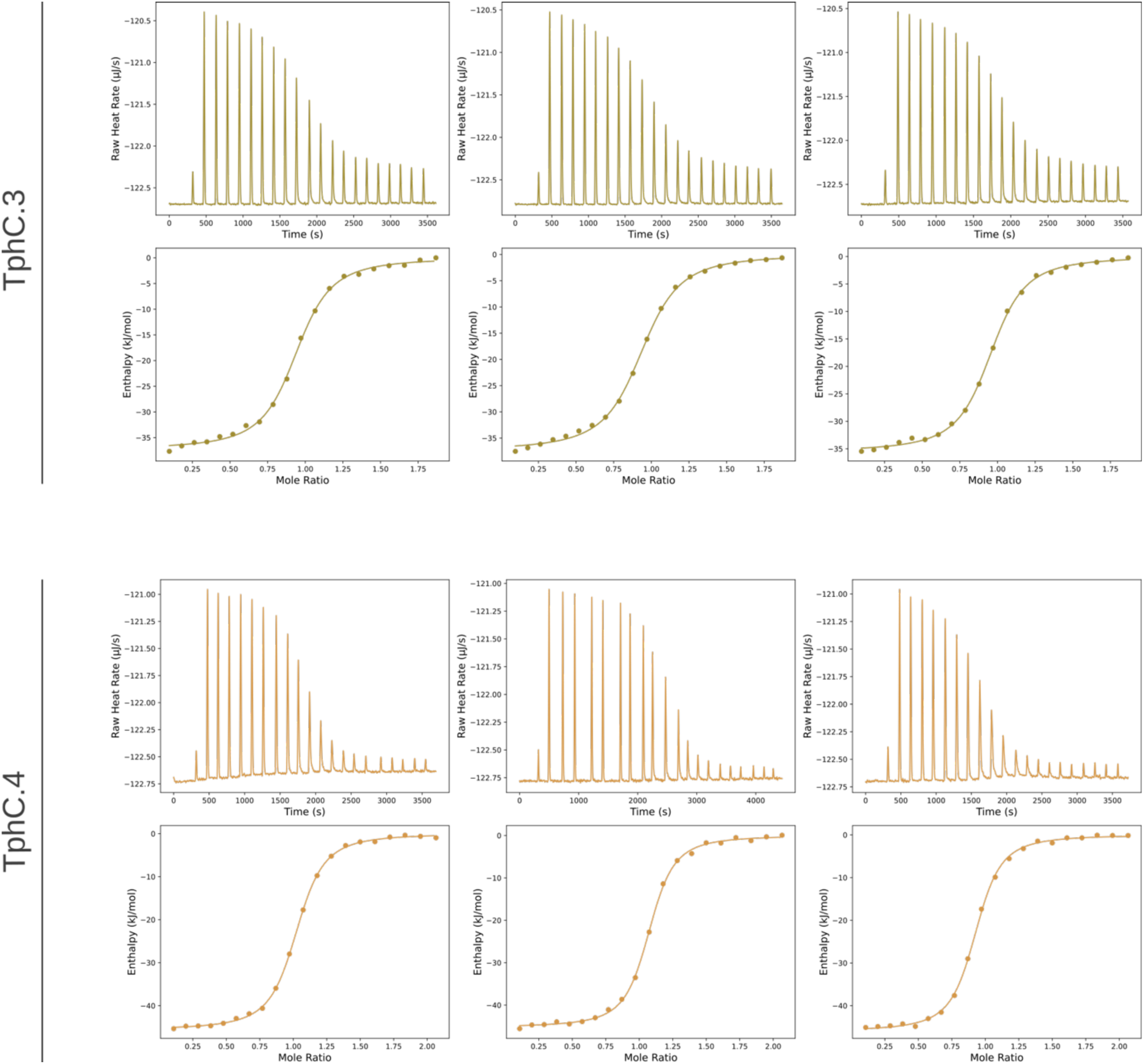
Thermograms and binding isotherms of TphC.0-4 and terephthalate in three technical replicates. Buffer-terephthalate control titrations were performed to obtain baseline values that were subtracted from the integrated heats of TphC.0-4.

**Supporting Figure 12.**
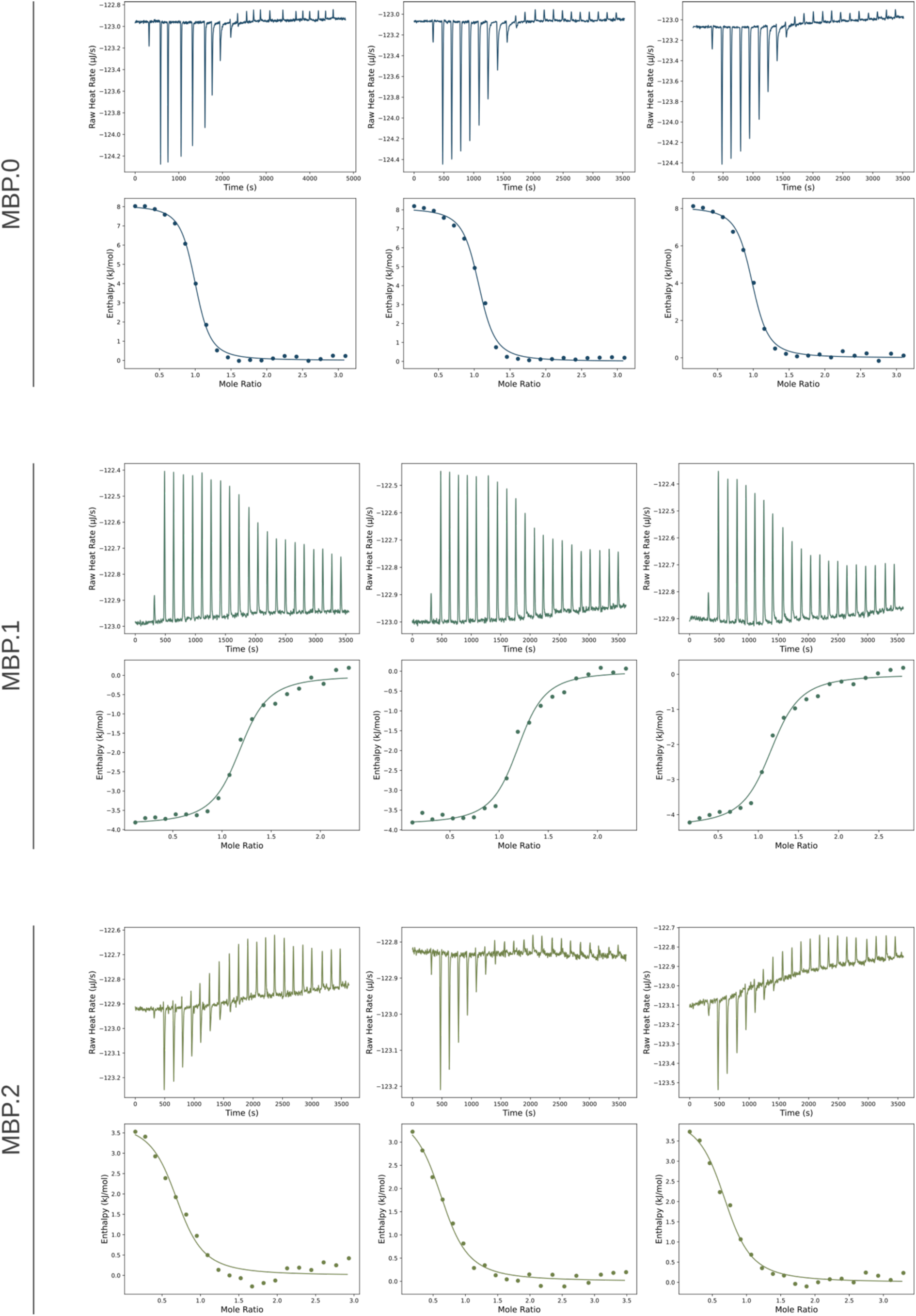
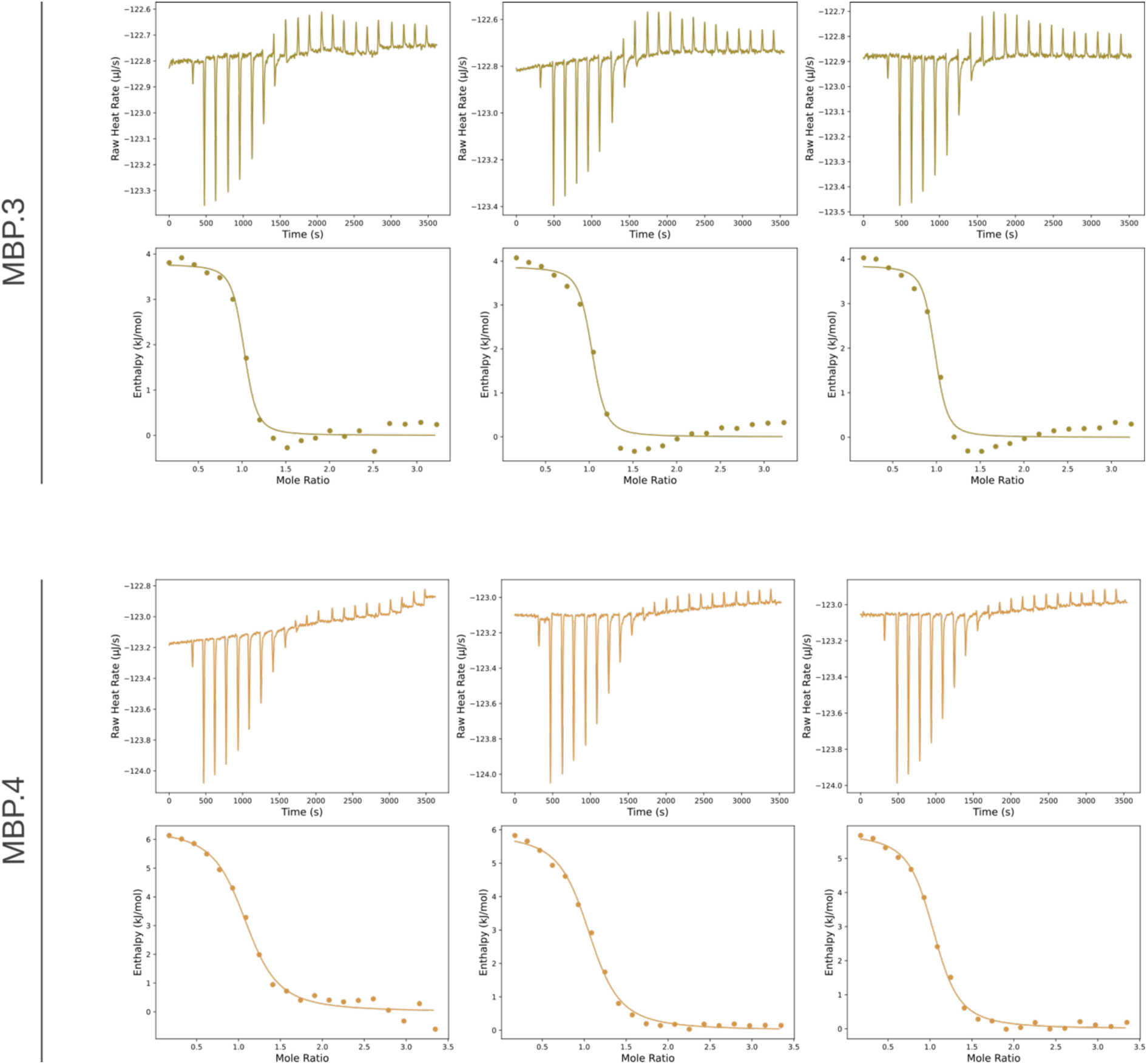
Thermograms and binding isotherms of MBP.0-4 and maltose in three technical replicates. Buffer-maltose control titrations were performed to obtain baseline values that were subtracted from the integrated heats of MBP.0-4.

**Supporting Figure 13.**
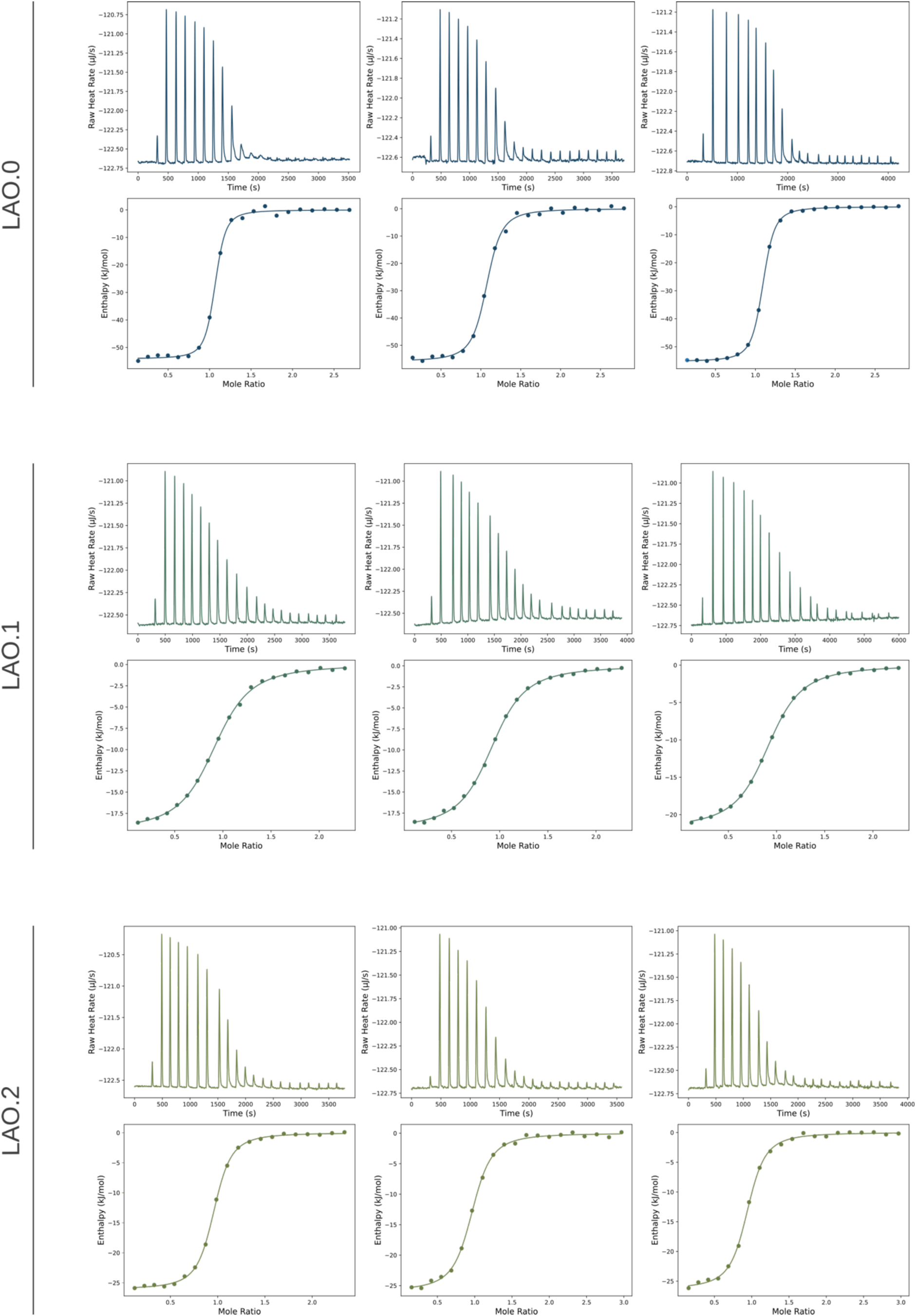
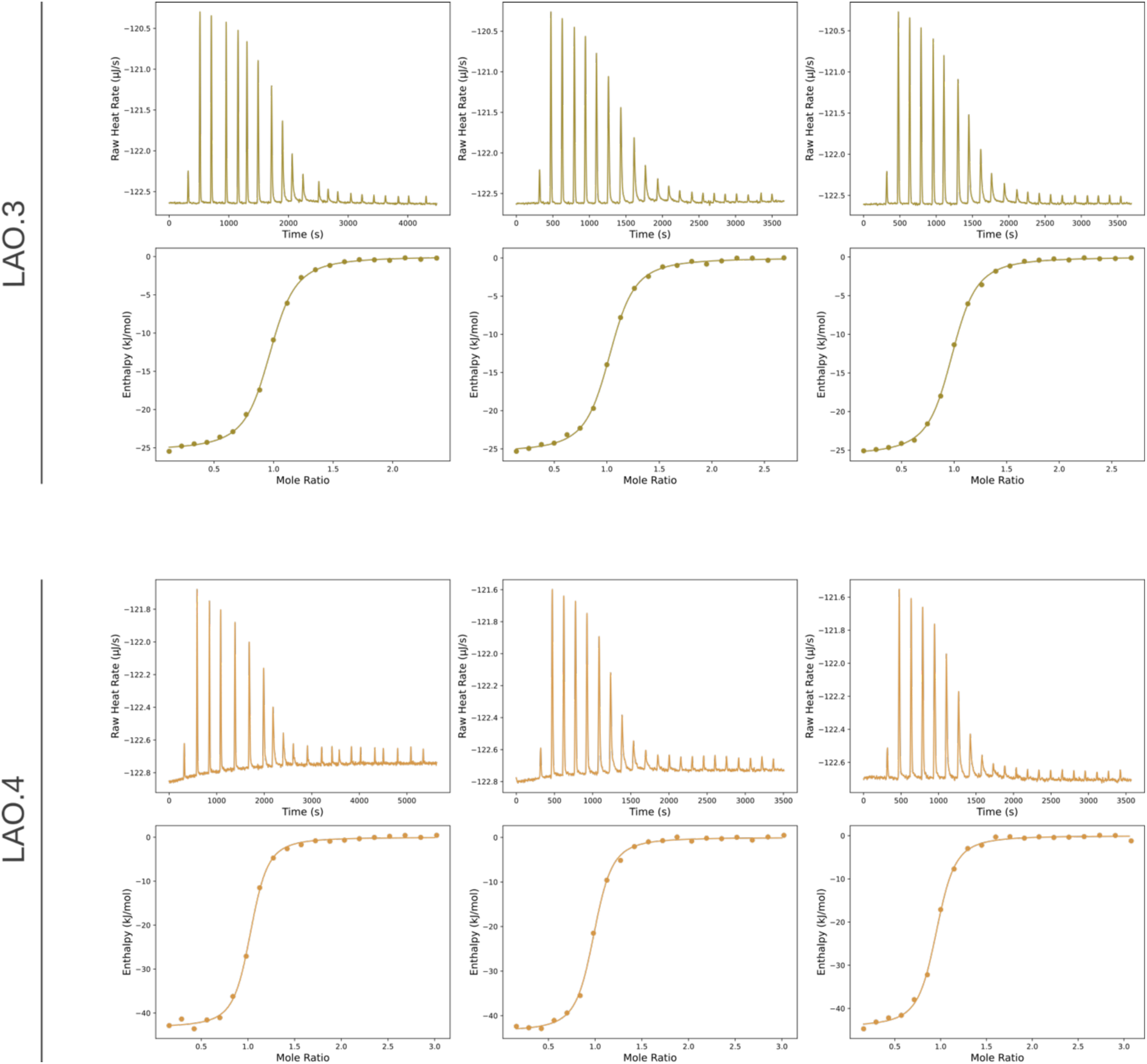
Thermograms and binding isotherms of LAO.0-4 and L-lysine in three technical replicates. Buffer-lysine control titrations were performed to obtain baseline values that were subtracted from the integrated heats of LAO.0-4.

